# Photoproximity labeling of c-Myc reveals SLK as a cancer specific co-regulator

**DOI:** 10.1101/2025.10.02.680136

**Authors:** Ryan R. Milione, Feifei Tong, Kelsey L. Wolfe, Sara B. Linker, Xinmeng Jasmine Mu, James V. Oakley, Andrew R. Nager, Michalina Janiszewska, Ciaran P. Seath

## Abstract

Transcription factors (TFs) have long been aspirational therapeutic targets for the treatment of diseases, as their dysregulation is a common mechanism for altered cell states. Despite this, many TFs implicated in disease have disordered structures and lack canonical binding pockets, rendering them non-trivial targets for small molecule-based therapies. Directly inhibiting TF function has proven difficult, but indirect inhibition by targeting the effector molecules that modulate TF function is a promising, yet underexplored, alternative approach. Here we report a strategy for capturing cancer-specific protein-protein interactions using context-dependent µMap photoproximity labeling. Using an intein-based method for catalyst conjugation in biochemically intact nuclei, we demonstrate that we can capture unique protein interactomes of c-Myc in healthy and cancerous prostate cell lines, and that these unique interactors can be mined to identify druggable vulnerabilities. We find that a cancer specific Myc interactor, SLK, selectively promotes c-Myc stabilization at the protein level, drives epithelial morphology, and is essential for tumorigenesis, validating it as a viable therapeutic target. Importantly, this stabilization is driven by a change in SLK splicing rather than expression at the protein or RNA levels. Furthermore, analysis of cancer patient data shows a strong correlation between this splice isoform and expression of c-Myc targets, suggesting this novel regulatory axis is operative across human cancer.

## Introduction

Transcription factors (TF) remain one of the most compelling yet elusive targets in cancer biology.^1^ They control gene expression programs that drive cancer progression, yet their disordered structures and lack of binding pockets have rendered direct inhibition challenging.^2^ TF activity depends on dynamic protein–DNA and protein–protein interactions, with these cistromic elements guiding localization and function.^3^ These interactions have presented opportunities for therapeutic disruption, although disease specificity is rarely considered, leading to clinical challenges as many TFs are also essential for normal tissue homeostasis.^4,5^ Furthermore, the oncogenic makeup of any cancer can create a distinct context for TF interactomes which can result in divergent responses to inhibition of a particular protein-protein interaction.^6,7^ Despite this, context dependent investigation of oncogenic TFs has yet to be realized.

The oncogenic transcription factor c-Myc controls the expression of thousands of genes associated with growth and is deregulated in over 40% of cancers.^8^ This master regulator of oncogenesis is characterized by its complex network of effector proteins that modulate its function^9^ and pharmacological interrogation of these interactors has been shown to be an effective method for treating c-Myc driven cancers in model systems.^8,10^ To identify co-regulating proteins, Penn and others have performed landmark studies investigating the interactome of c-Myc through a variety of methods, including co-immunoprecipitation and BioID proximity labeling (PL).^11–14^These methods have led to long lists of putative interactors that have been challenging to apply to the development of therapies as they broadly label transcriptional active regions of chromatin rather than direct c-Myc regulators. Furthermore, these approaches have never been applied contemporaneously to matched cancer cell lines to identify interactors specific to a single cancer phenotype or subtype.

We recently reported a nanoscale proximity labeling method (µMap) that bears a dramatically reduced labeling radius when compared to existing methods (4 nm vs 100 nm).^15,16^ This short radius has been shown to enable proximity labeling experiments that are sensitive to small changes in protein structure and environment.^15,17^ Furthermore, MacMillan and Muir have shown that µMap catalysts can be tracelessly incorporated onto histones using ultrafast split-intein splicing.^15^ Based upon these precedents, we sought to translate this method to c-Myc and extend this technology to capture cancer specific protein interactors. We reasoned that such a strategy would identify unique context dependent interactomes that can be mined to identify protein-protein interactions that modulate c-Myc at aberrant genetic loci that are only present in rapidly proliferating cells (Figure 1).

**Figure 1.**
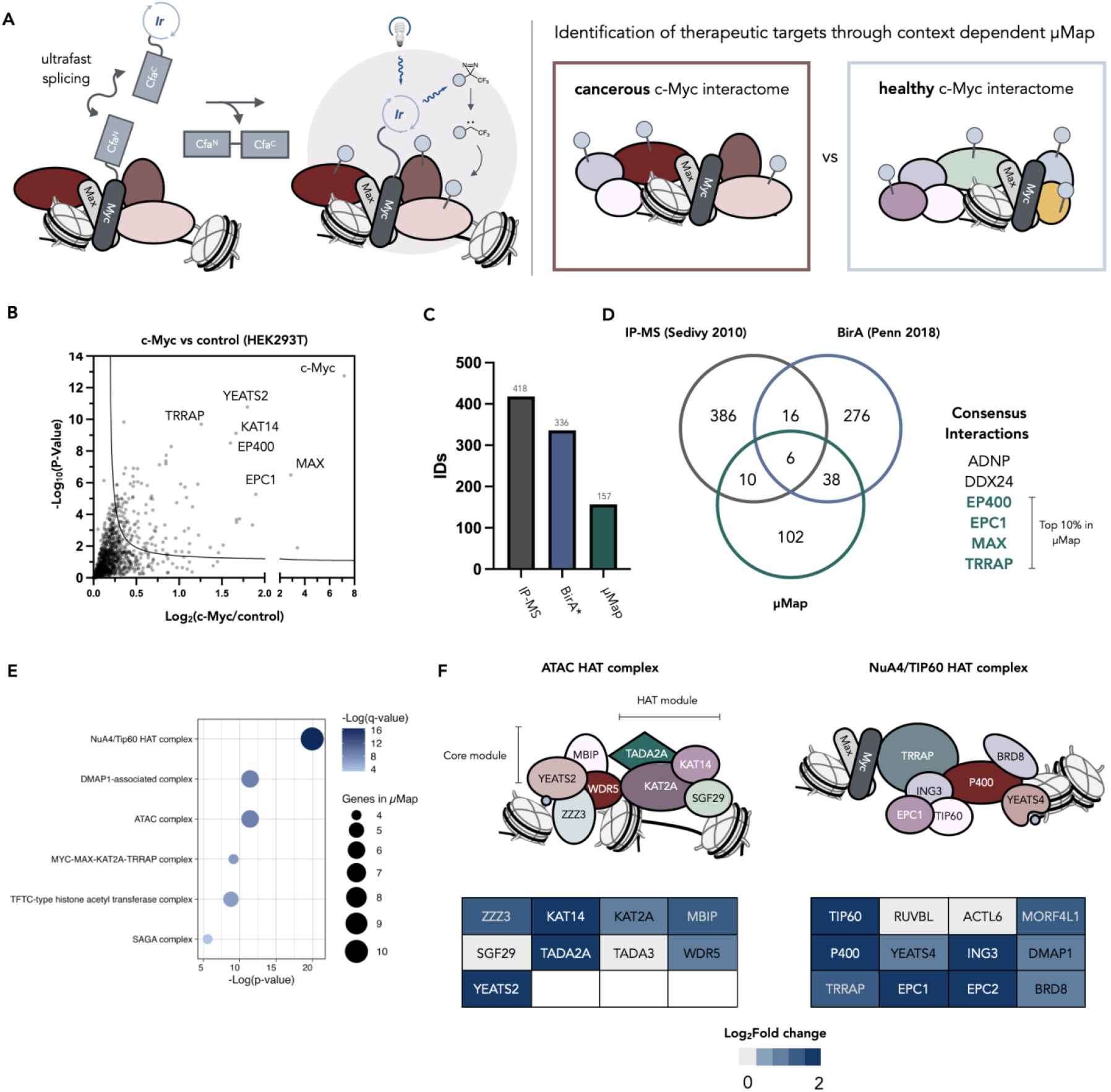
µMap identifies known c-Myc co-regulatory proteins in HEK293T cells. **a**, Cartoon schematic of µMap photoproximity labeling *in nucleo*. The iridium photocatalyst is conjugated to c-Myc via ultrafast split-intein splicing. After one minute of blue light irradiation, reactive diazirine probes bearing a biotin handle decompose to form reactive carbenes that crosslink with nearby proteins (∼4 nm radius). Biotinylated proteins are then enriched and characterized by mass spectrometry. Execution of this method in healthy and cancerous cell lines can reveal druggable cancer-specific c-Myc interactions. **b**, Mass spectrometry analysis after performing µMap in HEK293T cells transfected with c-Myc-CfaN. The bait protein (c-Myc), and known c-Myc co-regulatory proteins are among the most significantly enriched. **c, d**, When compared with other methods, µMap produces a refined list of c-Myc interacting proteins due to its reduced labeling radius. Six proteins are identified by all three methods, and four—MAX, EP400, TRRAP, EPC1—are among the top 10% enriched proteins by µMap. **e, f**, Protein complexes known to regulate c-Myc at chromatin are detected using µMap, including the NuA4/Tip60 and ATAC histone acetyltransferase (HAT) complexes.

## Results

### Technology development

We initiated our study by generating full length c-Myc constructs bearing a C-terminal intein tag (CfaN) flanked by FLAG and HA epitope tags for analytical convenience. Expression of this construct in HEK293T cells showed a single band at 75 kDa by western blot that could be stained with FLAG and HA antibodies, suggesting the fidelity of the construct is maintained upon expression (Fig. S1A,B). Next, we demonstrated the construct can splice with the CfaC peptides bearing either biotin or iridium handles. Incubation of biochemically intact nuclei with 1.0 µM CfaC-biotin/Ir for 40 minutes at 37 °C led to robust splicing, with approx. 95.5% conversion of the initial construct (Fig. S1B). Irradiation of spliced nuclei led to significant biotinylation and enrichment of c-Myc following streptavidin immunoprecipitation (Fig. S1D). Confocal microscopy of transfected cells showed clean nuclear localization of the construct, in line with the known function of c-Myc (Fig. S1C).

We next validated that our c-Myc construct remained functionally active and promoted the activation of Myc dependent genes. RNA-seq analysis showed similar changes in the expression of Myc controlled genes (HK2, SORD, UNG, HNRNPC) when compared to an unmodified c-Myc plasmid, suggesting our construct maintains transcriptional activity (Fig. S1E,F).

Confident in the validity of our PL system, we performed label free chemoproteomics using data independent analysis to capture the interactome of c-Myc using our µMap method in HEK293T cells. Our first experiment compared the c-Myc interactome to an identical protocol without transfection of the c-Myc-CfaN transgene. Each condition was performed with three biological replicates with two technical replicates per biological replicate to account for any inconsistencies in the protocol. For hit-determination, we established cut-offs of Log_2_(Fold change) ≥ 0.25 and FDR corrected P-value < 0.05, and hits were found in all positive replicates to be considered. We obtained 3,503 protein IDs with 3,291 IDs being found in >75% of all injections. Compartmental analysis of the identified proteins showed 61.7% of all IDs were localized to the nucleus, consistent with flow cytometry data from our nuclear isolation protocol (Fig. S2A,B). Principal component analysis (PCA) of all replicates showed a clear difference between untransfected conditions and c-Myc localized labeling (Fig. S2C).

We observed strong enrichment of the c-Myc bait protein (Log_2_Difference = 7.17, P-value = 1.78e^-13^), in addition to 156 other hits that meet significance (Fig. 1B). We next compared our list of interactors with several other c-Myc interactomes from the literature to look for enrichment of common c-Myc transcriptional co-regulators. We utilized a co-IP dataset from Sedivy et al. and a BioID proximity labeling dataset from Penn and co-workers.^11,13^ These datasets identified 418 and 336 hits respectively, with 22 genes identified in both datasets. Our method provided a more restricted interactome of 157 hits and showed clear enrichment of 6 of these 22 common interactors, four of which were found in our top 10% enriched genes (Fig. 1C,D). These four proteins— MAX, EP400, TRRAP, EPC1—are well characterized c-Myc interactors.^12,14,18–20^ Furthermore, 70% of our top 40 most enriched genes were annotated interactors of c-Myc (IntAct database) (Fig. S2D). GO analysis of the dataset showed significant enrichment of proteins associated with histone acetylation, transcription coregulator activity, and DNA-binding transcription factor binding (Fig. S2E,F). Specific analysis of our interactome showed significant enrichment of the ATAC and NuA4/TIP60 histone acetyltransferase complexes, known functional regulators of c-Myc (Fig. 1E,F).^19,21^ Finally, to probe the ability of our approach to measure subtle changes to the chromatin bound c-Myc interactome, we treated c-Myc-CfaN transfected HEK29T cells with vehicle or 1µM JQ-1 prior to photolabeling. In this experiment we observed clean enrichment of BRD2/3/4 only in the untreated samples, showing that JQ-1 blocks BET binding to chromatin and consequently disrupts the Myc-BET axis (Fig. S2G).^22^

### c-Myc interactome in prostate cell lines

Androgen receptor (AR) negative prostate cancer is an aggressive form of prostate cancer that no longer responds to frontline treatment options such as androgen deprivation therapy, resulting in poor prognoses.^23^ Expression of AR in these tumors is often low or null, and growth is promoted by c-Myc, with few treatment options available for intervention.^24,25^ We questioned whether our proximity labeling approach would be sensitive enough to identify cancer specific transcriptional co-activators that may lead to c-Myc activation and subsequent growth and metastasis. To identify factors that may contribute to this effect, we compared the c-Myc interactome in three prostate cell lines that represent healthy prostate cells (WPMY-1), AR-negative prostate cancer (PC-3), and AR-positive prostate cancer (LNCaP). In both AR- (PC-3) and AR+ (LNCaP) cell lines, c-Myc is primarily stabilized at the protein level rather than being transcriptionally upregulated, suggesting changes in protein structure or interactome may promote cancer progression through increased c-Myc activation (Fig. S3A,B).

Comparison of the three interactomes revealed several common co-regulating proteins such as MAX, MORF4L1, MORF4L2, and CCAR2, suggesting that these interactions are critical for c-Myc function in both healthy and cancerous contexts (Fig. S3C,D). We observed significant differences in the enrichment of proteins associated with protein degradation (proteosome, E3 ligase machinery) across the three cell lines, with highest enrichment in WPMY-1 cells and lowest in LNCaP, which correlated with the half-life of c-Myc (Fig. S3E,F). As c-Myc degradation has been implicated to be controlled by phosphorylation status^26–29^, we compiled all enriched kinases across all cell lines tested and found a panel of unique kinase interactors in each along with several c-Myc associated kinases that were enriched in both cancer lines but not in the healthy prostate line (Fig. S3G). These include CDK1, CDK9, CDK11B, and PLK1, all of which are critical regulators of cellular growth. To identify cancer specific co-regulators that modulate cellular phenotypes we cross-analyzed the enriched proteome with DepMap RNAi screens to identify c-Myc interactors that are important for cell survival. In PC-3, we identified a single druggable protein that met our filters, STE20-like kinase SLK (Fig. 2B,C).

**Figure 2.**
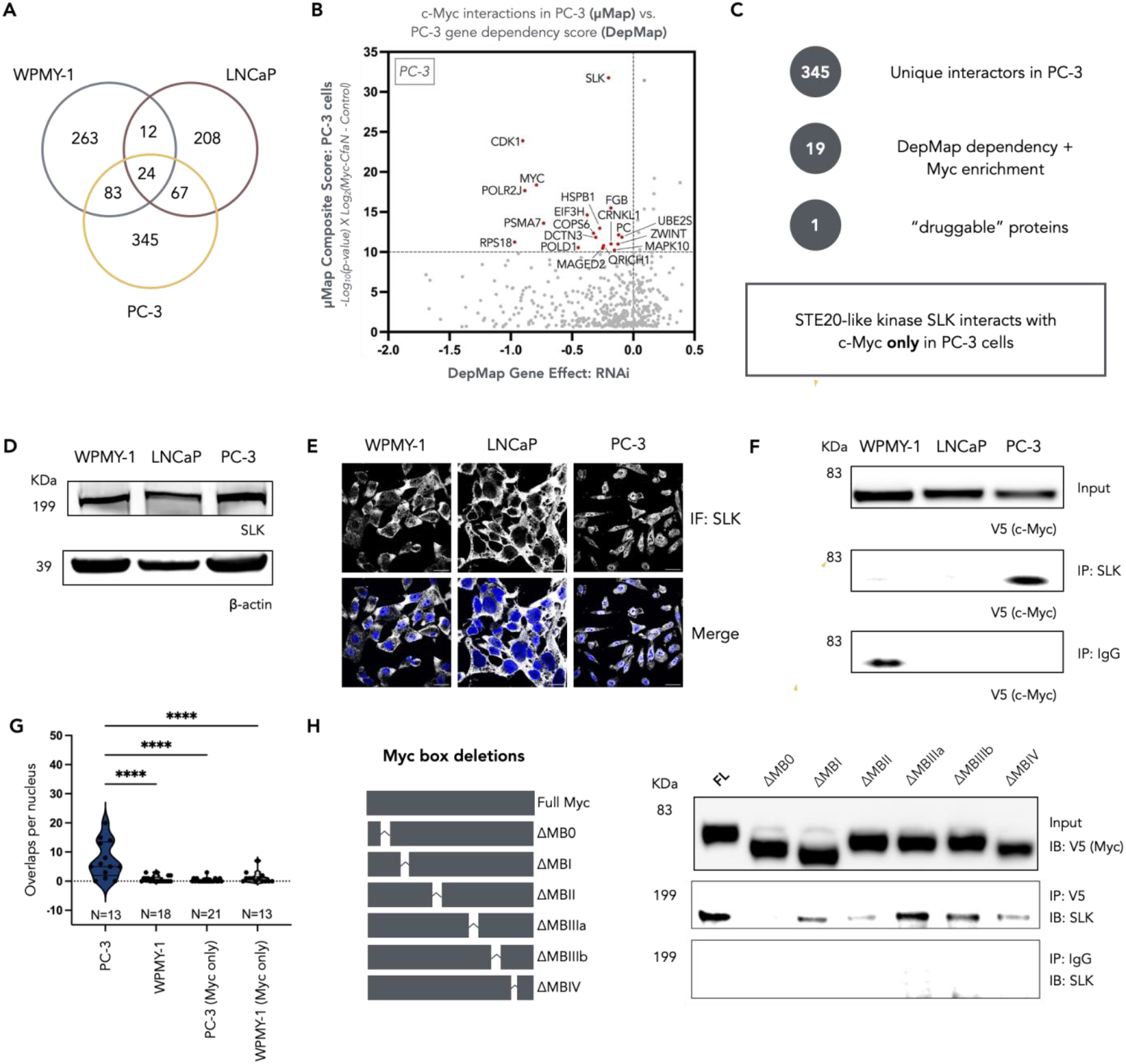
µMap identifies a novel interaction between SLK and c-Myc that is cancer specific. **a**, Venn diagram showing c-Myc interactions enriched by µMap in healthy prostate cells (WPMY-1), and metastatic prostate cancer cells (LNCaP, PC-3). **b, c**, All c-Myc interactions enriched by µMap in PC-3 cells are plotted against their corresponding DepMap dependencies. STE20 like kinase (SLK) only interacts with c-Myc in PC-3 cells and is the most enriched protein in µMap (composite score = 31.76) **d**, Endogenous expression of SLK across the three cell lines is uniform. **e**, Immunofluorescent staining of SLK suggests the kinase is primarily localized in the cytoplasm in WPMY-1 and LNCaP cells but is localized in both the nucleus and cytoplasm in PC-3 cells. **f**, Co-immunoprecipitation of SLK suggests c-Myc only binds SLK in PC-3 cells. Scale bar = 30µm (PC-3, WPMY-1); scale bar = 20µm (LNCaP). **g**, Immunofluorescent staining of c-Myc and SLK with rolling circle amplification (Molboolean™ assay) suggests that c-Myc and SLK co-localize in the nucleus of PC-3 cells, but not in the nucleus of WPMY-1 cells. **h**, Deletion of Myc-box 0 ablates recovery of SLK after c-Myc immunoprecipitation. ^*^P < 0.05, ^**^P < 0.01, ^***^P < 0.001, ^****^P < 0.0001; ns, not significant.

### SLK interacts with c-Myc only in PC-3 cells

SLK is involved in focal adhesion and microtubule organization, but very little research has been dedicated to its role in cancer, and it has never been associated physically or genetically to c-Myc.^30–32^ We assessed the expression levels of SLK in all three cell lines by western blotting, showing similar levels of the protein with no increase observed in PC-3 cells that would account for differential enrichment (Fig. 2D). Confocal microscopy of SLK confirmed the expression levels in all cell lines yet showed an altered distribution only in PC-3 cells, which exhibited a significant increase in nuclear SLK (Fig. 2E). Co-IP experiments showed a robust Myc-SLK interaction in PC-3 but not in LNCaP or WPMY-1 cells (Fig. 2F). Additionally, recovery of Myc protein after IP of SLK in PC-3 cells was strengthened following nuclease treatment, suggesting the interaction between SLK and c-Myc is not dependent on DNA scaffolding (Fig. S4A). MolBoolean analysis of the two proteins also showed significant colocalization within the nucleus that is unique to PC-3 cells (Fig. 2G, S4B). Co-IP with Myc-box deletions showed this interaction was dependent on MB0, the transactivation domain of c-Myc (Fig. 2H).

Two different splice isoforms are known for SLK that differ through inclusion of exon 13, leading to a change of 31 amino acids that is thought to impact protein dimerization.^33–35^ ESRP1 and ESRP2 promote exon 13 inclusion, generating the long form (SLK-L),^34,36,37^ and RBFOX2 promotes exon 13 skipping, generating the short form (SLK-S).^33,38^ Some evidence has suggested that isoform selectivity leads to altered proliferation in cancer^38,39^ and cellular localization.^40^ Recent research shows that nuclear localization of SLK is dependent on the inclusion of exon 13, which contains a nuclear localization sequence (NLS).^40^ To assess if the change in SLK localization and c-Myc-binding may be attributed to differential expression of SLK isoforms, we performed splicing analysis on each cell line to assess altered splice isoforms in the SLK transcript. These data showed inclusion of exon 13 of SLK only in PC-3 cells, which correlated with increased expression of ESRP1/2 and reduced expression of RBFOX2 (Fig. 3A-C). We then evaluated publicly available RNA sequencing data for expression of the two SLK isoforms in cancer and healthy samples.^41,42^ In prostate cancer samples, the SLK-L isoform represents ∼75% of SLK transcripts, but in normal prostate samples SLK-S is the dominant transcript (Fig. 3D). This trend held true when including all samples and tissue types in the analysis, suggesting the SLK-L isoform is oncogenic (Fig. 3E). Furthermore, expression of SLK-L, not SLK-S, correlates with expression of c-Myc target genes in prostate cancer, supporting a mechanism where this isoform drives oncogenesis by increasing c-Myc activity (Fig. 3F, Fig. S4C-E).

**Figure 3.**
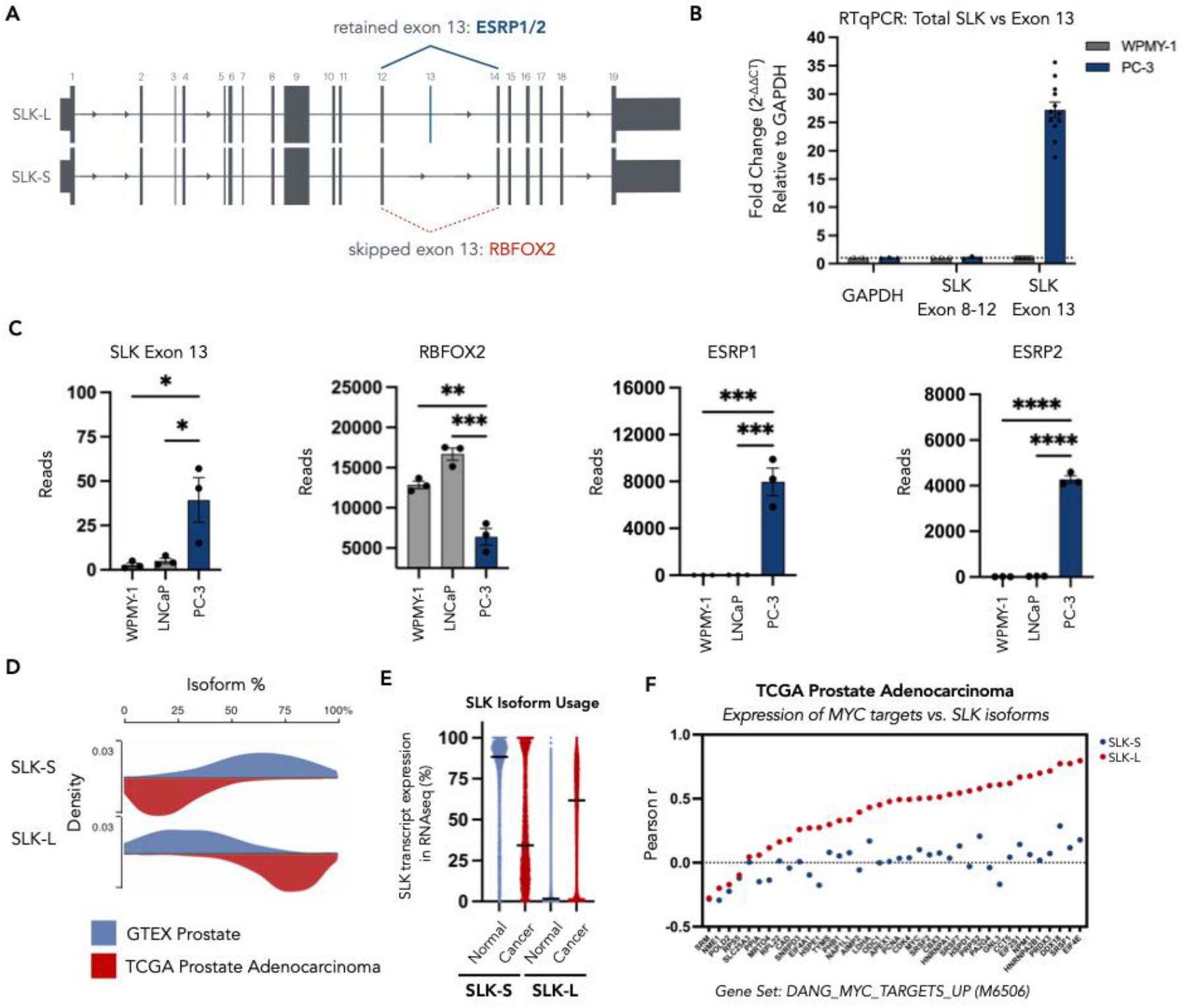
PC-3 cells express the long splice isoform of SLK, which can localize to the nucleus. **a**, SLK has two alternative splice isoforms, SLK-L and SLK-S, differing only by the presence of exon 13, which is 93 base pairs in length. Retention of exon 13 is governed by epithelial splicing regulatory proteins 1 and 2 (ESRP1/2), and exon 13 skipping is regulated by RNA-binding fox-1 homolog 2 (RBFOX2). **b**, RTqPCR analysis using primer pairs spanning SLK exon 8-12 (total SLK) or primer pairs specific for SLK exon 13 (SLK-L), suggests that PC-3 cells have ∼25-fold more SLK-L mRNA transcripts than WPMY-1 cells. **c**, Total RNA sequencing reads for SLK exon 13, RBFOX2, ESRP1, and ESRP2. PC-3 cells have increased expression of SLK-L and ESRP1/2 mRNA, and decreased expression of RBFOX2 mRNA compared to WPMY-1 and LNCaP cells. **d-f**, Analysis of RNA sequencing data from the TCGA TARGET GTEX study on the UCSC Xena database (xenabrowser.net).^41,42^ **d**, SLK-S is the major isoform present in normal prostate samples (GTEX), while SLK-L is the major isoform present in prostate adenocarcinoma (TCGA). **e**, Across all samples in the study, SLK-S is the major isoform in normal samples (GTEX, N = 7,862), but SLK-L is the major isoform in cancerous samples (TCGA, N = 10,535). **f**, Expression of c-Myc target genes in prostate adenocarcinoma samples correlates with expression of the SLK-L isoform, but not SLK-S expression. ^*^P < 0.05, ^**^P < 0.01, ^***^P < 0.001, ^****^P < 0.0001; ns, not significant.

### SLK determines c-Myc stability in PC-3 cells

We next investigated the role SLK plays in c-Myc based transcription. CRISPR-KO of SLK in PC-3 cells had a dramatic effect on c-Myc protein levels, reducing c-Myc to ∼20% of WT PC-3 cells (Fig. 4A). RT-qPCR analysis showed this change was for the most part at the protein level rather than a loss of transcription of the mRNA, which was confirmed by cycloheximide chase which showed a 2.3-fold reduction in half-life in SLK-KO PC-3 cells (138 min to 60 min) (Fig. 4B, C).We next investigated whether the significant decrease in c-Myc levels led to an associated decrease in proliferation and migration. ATP-based proliferation assays showed diminished cell division in the SLK-KO cells, with doubling times increasing from 52h to 79h (Fig. 4D). A scratch assay to simulate wound healing showed a similar trend where SLK-KO and pharmacological inhibition repressed wound closure (Fig. 4E). Wound healing and proliferation were only minimally affected by SLK-KO in WPMY-1, suggesting a cancer specific effect of SLK (Fig. 4D, Fig. S4F). Additionally, cell cycle analysis using propidium iodide showed a readjustment of the cell cycle, with a loss of S phase cells and a corresponding increase in G2 phase (Fig. 4F, Fig. S4G). These data are consistent with prior reports of transcriptional silencing of c-Myc using siRNA in PC-3, suggesting these observed phenotypes arise through Myc depletion.^43,44^ Furthermore, xenograft models of PC-3 and SLK-KO cells showed complete inhibition of tumorigenesis in SLK-KO, demonstrating the importance of SLK for cancer growth (Fig. 4G).

**Figure 4.**
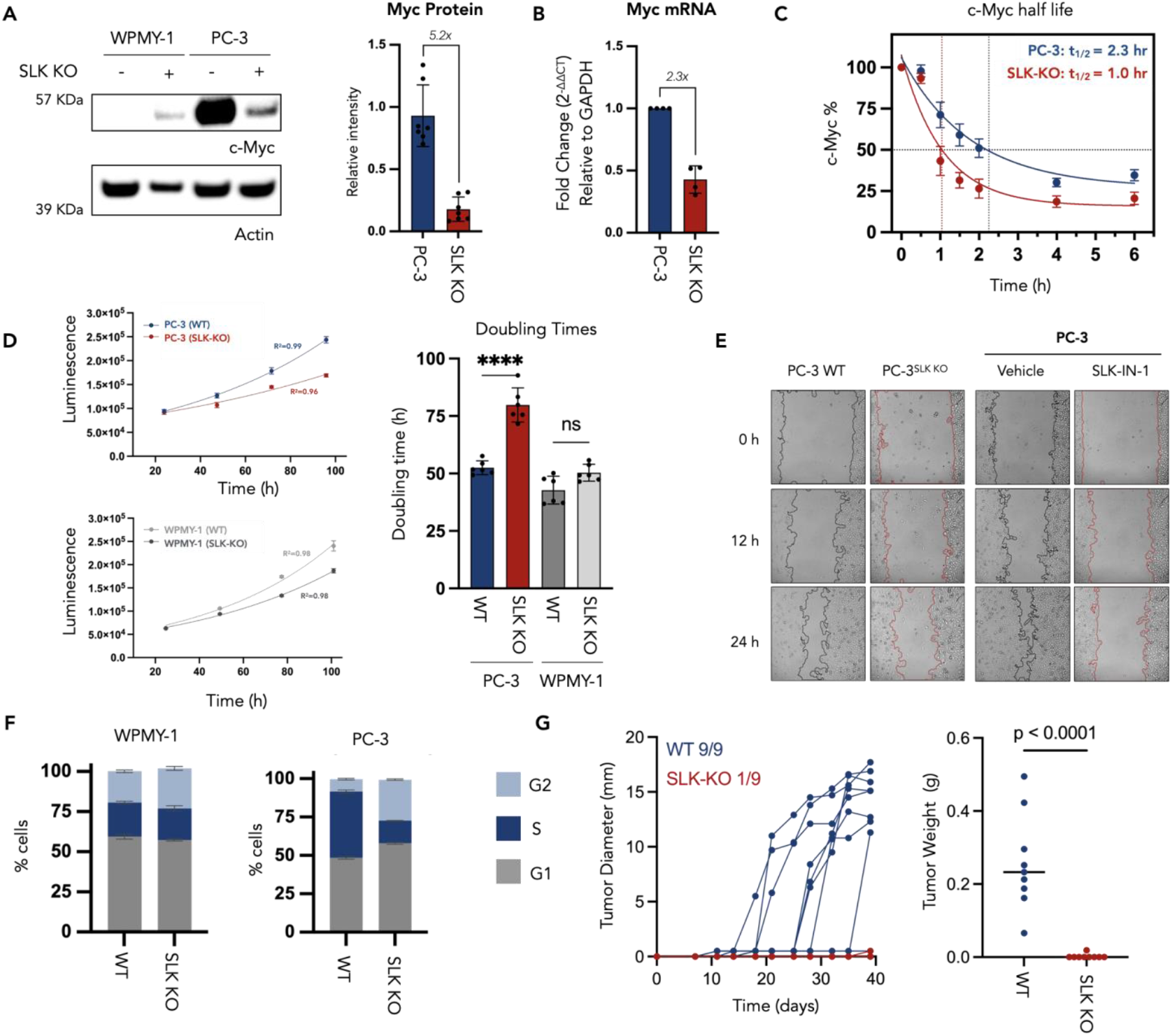
SLK stabilizes c-Myc protein and promotes growth and migration of PC-3 cells only. **a**, CRISPR knockout of SLK in PC-3 cells reduces c-Myc protein expression by 5.2-fold, while knockout of SLK in WPMY-1 cells leads to a modest increase in c-Myc protein. **b**, RTqPCR analysis in PC-3 cells suggests c-Myc mRNA is reduced 2.3-fold after SLK knockout. **c**, Cycloheximide chase assays suggest c-Myc protein half-life is reduced by 1.3 hours in the SLK knockout PC-3 cells. **d**, Doubling times measured using the ATP-based CellTiter-Glo 2.0 reagent. SLK knockout increases the doubling time of PC-3 cells by 27 hours and the doubling time of WPMY-1 by only 8 hours. **e**, 24-hour scratch assays suggest that PC-3 cell migration is inhibited by SLK knockout and a commercially available SLK inhibitor (SLK/STK10-IN-1, 10µM). **f**, Cell cycle analysis with propidium iodide. SLK knockout in PC-3 cells reduces the percentage of cells in S-phase and leads to G2 stalling, while WPMY-1 cell cycle is minimally affected by SLK knockout. **e**, *In vivo* experiment with wild-type and SLK knockout PC-3 cells. Flank injection of wild-type PC-3 cells leads to rapid tumor formation, while SLK knockout PC-3 cells fail to form tumors. ^*^P < 0.05, ^**^P < 0.01, ^***^P < 0.001, ^****^P < 0.0001; ns, not significant.

To better understand these phenotypes, we compared the proteomes and phosphoproteomes of the WT and SLK-KO cells. We observed significant remodeling of the cellular proteome. Global proteomics demonstrated a significant loss of epithelial markers (EPCAM, EGFR) and proteins associated with growth and chromatin regulation (BRD4, MKI67, AURKB, MCM complex, CDCA8, PLK1, CTNNB1) (Fig. S5A-C). ATG5, ATG12, ATG16L1, GABARAPL2, WIPI2, OPTN), and the interferon response (MX1, MX2, IFIT1, IFIT2, IFIT3, IFIT5, IFI44, IFI44L, IFI16, ISG20) (Fig. S5D-F). We validated a selection of the most enriched proteins by either western blot, immunofluorescence, and flow cytometry, confirming the accuracy of our analysis (Fig 5C,D,F, Fig. S5G-H, Fig. S6A-C). The phosphoproteome of the wild-type PC-3 cells were enriched in phosphosites related to the cell cycle and chromatin organization, whereas SLK-KO led to enrichment of mRNA metabolism, splicing, and GTP signaling (Fig. S7A,B). Additionally, Myc and EGFR activity were the most differentially enriched pathways in wild-type PC-3 cells, consistent with SLK-KO repressing c-Myc based transcription (Fig. 5B). ATM was the most active kinase in SLK-KO based on substrate analysis, suggesting a loss of chromatin integrity and DNA damage (Fig. S7C,D). The Myc phosphodegron (T58, S62) associated with its proteasomal degradation was enriched in the SLK-KO cell line, supporting our hypothesis that SLK binding inhibits Myc phosphorylation at T58 and prevents degradation of Myc by the proteasome (Fig. S7D,E). In addition to a loss of Myc expression, we observed significant morphological changes to the SLK deficient PC-3 cells over time (Fig S8B). Quantification of nuclear size at early passages showed a significant increase in SLK-KO compared to WT PC-3 (Fig. 5D, Fig. S8A). In agreement with our analysis of the phosphoproteome, we observed an increase in phospho-ATM (Ser1981) in SLK-KO, a marker for the DNA damage response (Fig. 5E). RT-qPCR analysis of markers of the epithelial to mesenchymal transition (SLUG, SNAIL, ZEB1) showed positive regulators of EMT are transcriptionally upregulated after SLK-KO (Fig. 5F, Fig. S6D). Taken together, these data suggest a mechanism where SLK-KO leads to loss of Myc stabilization, driving an epithelial to mesenchymal transition, subsequent loss of chromatin integrity, inhibition of autophagy, and activation of the interferon response.

**Figure 5.**
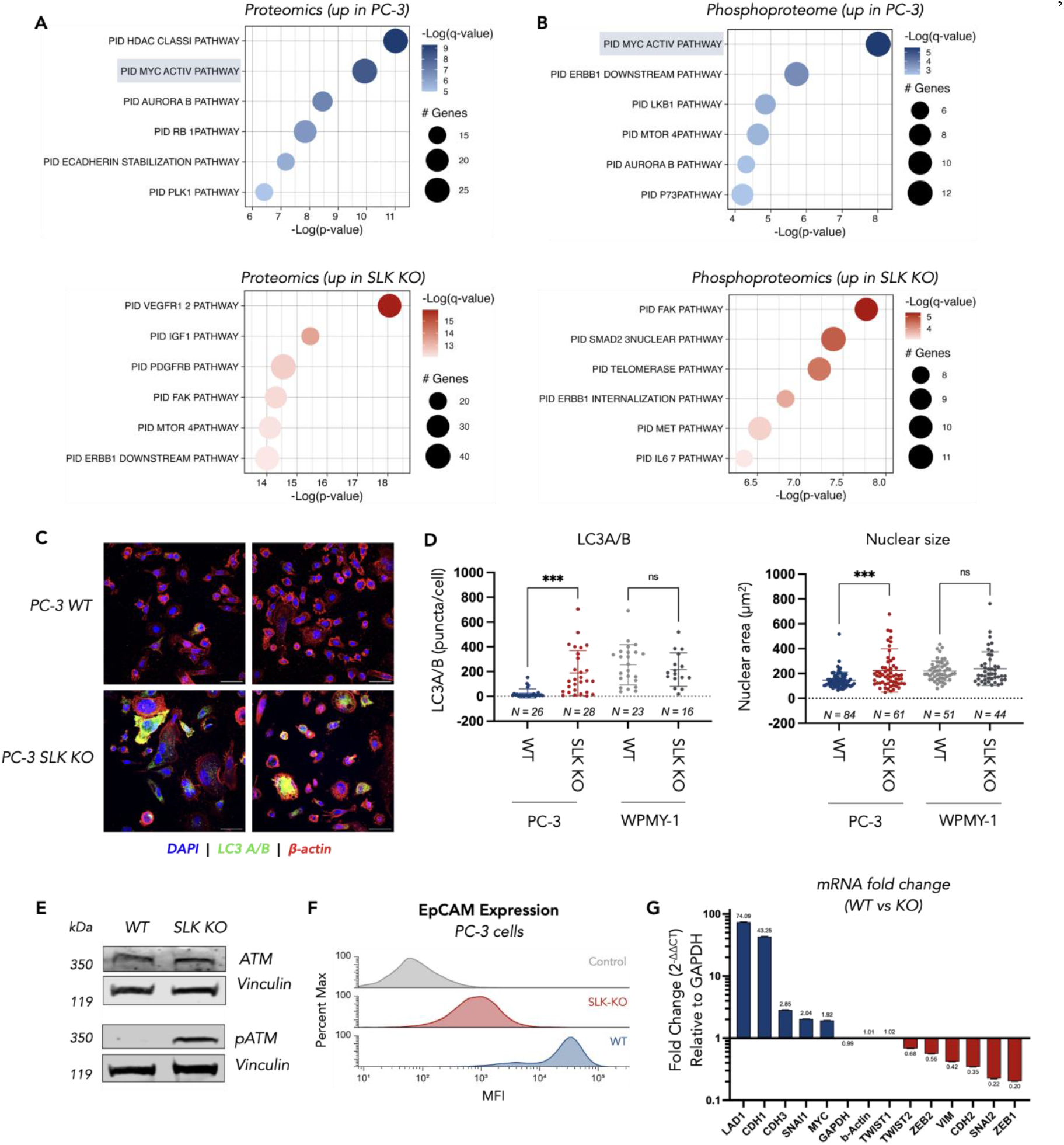
SLK regulates autophagy, maintains chromatin integrity, and promotes an epithelial phenotype in PC-3 cells. **a**, Canonical pathways upregulated in the global proteome of wild type and SLK knockout PC-3 cells. **b**, Canonical pathways upregulated in the phosphoproteome of wild type and SLK knockout PC-3 cells. **c**, Immunofluorescent staining of LC3 A/B suggests LC3 A/B proteins are upregulated in SLK knockout PC-3 cells. Scale bar = 20 µm **d**, Quantification of LC3 A/B signal and nuclear area (DAPI) in confocal microscopy images. LC3 A/B expression and nuclear area increase in the SLK knockout PC-3 cells, but there is no significant difference in the SLK knockout WPMY-1 cells. **e**, SLK knockout increases phosphorylated ATM in PC-3 cells, a marker of DNA damage. **f**, Analysis of cell surface EpCAM expression by flow cytometry suggests EpCAM expression is reduced in SLK knockout PC-3 cells. **g**, RTqPCR analysis of genes that dictate epithelial and mesenchymal cell states. Wild type PC-3 cells express markers of epithelial cells such as LAD1 and CDH1. SLK knockout PC-3 cells express markers of mesenchymal cells, such as ZEB1 and SNAI2. ^*^P < 0.05, ^**^P < 0.01, ^***^P < 0.001, ^****^P < 0.0001; ns, not significant.

## Discussion

In summary, we have developed a photocatalytic proximity labeling strategy for uncovering druggable, cancer specific interactions at c-Myc. The short range diazirine activation mechanism allows the collection of highly precise interactomics data that facilitate sorting with the DepMap database to identify drivers of oncogenic c-Myc signaling. In these experiments, we observe a specific splice variant of SLK which drives and nuclear localization and an interaction with c-Myc which promotes its stabilization at the protein level. Knockout of SLK impacts cellular growth in vitro and abolishes tumor formation in a xenograft model providing evidence that SLK inhibition or degradation may be a viable strategy for therapeutic intervention. Importantly, these cellular effects are specific, with SLK knockout having minimal impact on healthy prostate cells and an AR+ prostate cancer cell line. Analysis of healthy and cancerous patient samples shows that the SLK-Long isoform is broadly co-opted by cancer, and its presence correlates with expression of c-Myc targets, providing evidence for a mechanism where SLK stabilizes c-Myc and drives oncogenic transformation and proliferation beyond the prostate cell lines described in this study. In future, we hope to stratify tumor tissue by nuclear SLK staining to provide personalized cancer treatments for Myc-driven cancers by intervention through the SLK-Myc axis. Importantly, as SLK expression at the protein and RNA level is unchanged between healthy and cancerous contexts, identification of this important regulatory axis would be impossible without high resolution subcellular technologies such as µMap.

Thus, our results suggest that cell-context–specific protein–protein interactions may be critical for identifying novel therapeutic targets, stratifying patients in clinical settings, and ultimately reducing attrition in drug development while enabling more effective, personalized therapies. As transcription is generally templated by complex cistromes, we anticipate that this strategy will be applicable to diverse transcriptional proteins in various disease states, opening therapeutic avenues for yet-to-be identified co-regulators.

## Data Availability

All relevant data are included in the manuscript and supplementary information. Mass spectrometry data files will be uploaded to the MassIVE proteomics database. RNA sequencing data files have been uploaded to the GEO database (GSE307813).

## Acknowledgements

Research reported in this publication was supported by the Office of The Director, of the National Institutes of Health under Award Number S10OD036363 and the National Institute of General Medical Sciences of the National Institutes of Health (R35GM150765). We also acknowledge the Wertheim UF-Scripps for start-up funds. The content is solely the responsibility of the authors and does not necessarily represent the official views of the National Institutes of Health. The authors thank George Tsaprailis and Catherina Scharager Tapia at the UF-Scripps Proteomics Facility, Robert M. Witwicki, Li Pan, and Marlene L. Biller at the UF-Scripps Genomics Facility, and Gogce C. Cryen and Alexander Trouern-Trend at the Bioinformatics and Statistics Core.

The analyses of publicly available RNA sequencing data in this manuscript are based upon data generated by the TCGA Research Network (https://www.cancer.gov/tcga), the Therapeutically Applicable Research to Generate Effective Treatments (TARGET) initiative (phs000218; https://www.cancer.gov/ccg/research/genome-sequencing/target), and The Genotype-Tissue Expression (GTEx) Project (supported by the Common Fund of the Office of the Director of the National Institutes of Health, and by NCI, NHGRI, NHLBI, NIDA, NIMH, and NINDS). The data used for the analyses described in this manuscript were obtained from the UCSC Toil RNAseq Recompute Compendium.^41,42^

## Author Contributions

CPS conceived the work. CPS, RRM, FT, KLW, MJ, designed and executed the experiments. XPS, RRM, FT, KLW, MJ, SL, JVO, ARN, XJM analyzed the data, CPS, RRM, MJ, JVO prepared this manuscript.

## Author Information

The authors declare no competing financial interests.

