## Supplementary Figures and data for "Photoproximity labeling of c-Myc reveals SLK as a cancer specific co-regulator"

The PDF file includes:

Figs. S1 to S8
Materials and Methods
References

### Extended Data Figures

#### Extended Data Figure 1 – Validation of the c-Myc-CfaN transgene

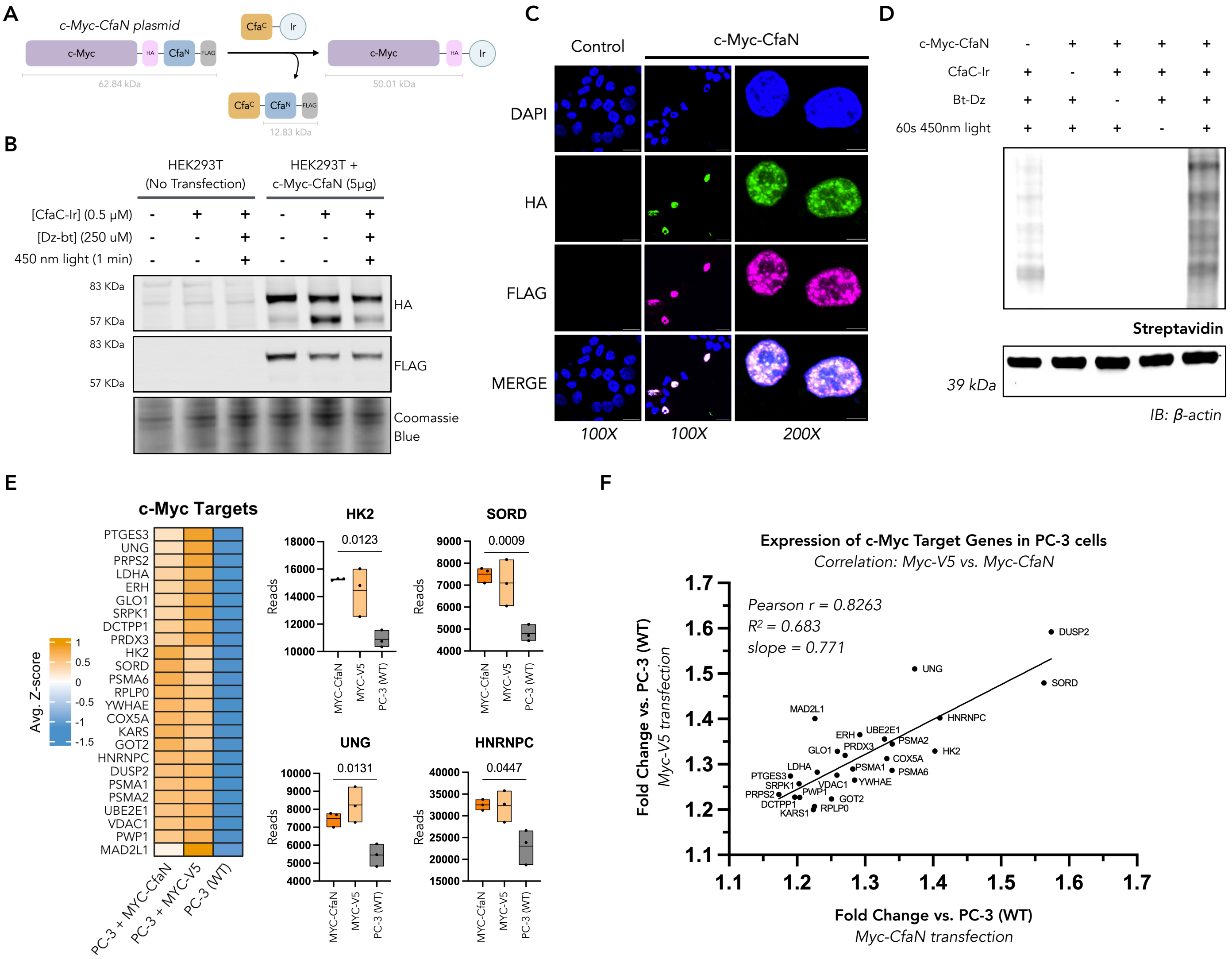


**Extended Data Figure 1. A)** The µMap iridium photocatalyst is conjugated onto c-Myc via ultrafast split-intein splicing. **B)** Western blot of nuclear lysates of wild-type or c-Myc-CfaN transfected HEK293T cells. Successful incorporation of the iridum photocatalyst is visualized by HA and FLAG staining. In the HA stain, the appearance of a lower band at 62 KDa after treating the nuclei with 0.5 µM CfaC-Ir at 37°C for 40 min suggests the full intein + FLAG (12.83 KDa) has been spliced out, and incorporation of the iridium photocatalyst on to c-Myc has occurred. Similarly, the intensity of the FLAG staining after CfaC-Ir treatment is reduced, further indicating splicing has occurred. **C)** The c-Myc-CfaN transgene localizes to the nucleus in LNCaP cells. Scale bar = 20µm (100X), 5µm (200X). **D)** All four components of µMap are required for sufficient labeling. Omission of the c-Myc-CfaN transgene, CfaC-Ir treatment, diazirine-PEG3-biotin (Bt-Dz) probes, or 60 seconds of 450nm light reduces total biotin labeling. **E, F)** The c-Myc-CfaN regulates Myc target genes. Total RNA sequencing suggests Myc target genes from HALLMARK_MYC_TARGETS_V1 and HALLMARK_MYC_TARGETS_V2[1] are upregulated in PC-3 cells transfected with c-Myc-V5-6xHis (control) or c-Myc-CfaN (µMap), when compared with wild-type PC-3 cells. For the Myc target genes shown in E, the relative expression after transfection of c-Myc-V5-6xHis (no intein) correlates with the relative expression after transfection of c-Myc-CfaN (bearing 12.5 KDa split-intein on the C-terminus), suggesting the fusion of the split-intein onto c-Myc does not inhibit interactions with co-regulatory proteins that facilitate transcriptional regulation of Myc target genes.

### Extended Data Figure 2 – µMap captures known c-Myc interactions in HEK293T cells

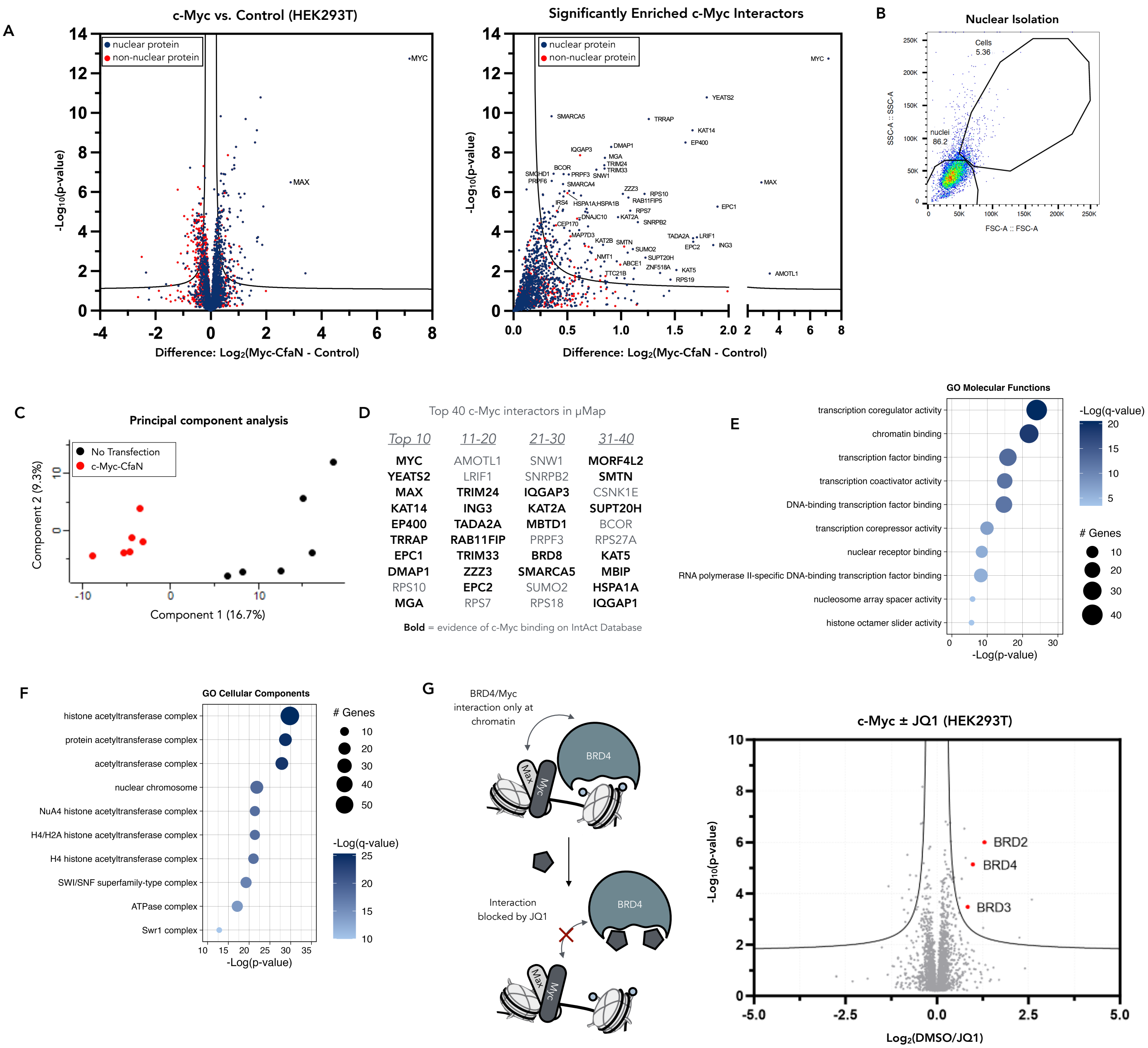


**Extended Data Figure 2. A)** (Left) A volcano plot of all proteins in mass spectrometry data after µMap in HEK293T cells. Nuclear proteins (navy) are preferentially enriched in the cells transfected with c-Myc-CfaN. (Right) Zoomed volcano plot showing significantly enriched c-Myc-proximal proteins are mainly nuclear. **B)** Flow cytometry suggests biochemically intact nuclei represent ~85% of the remaining population after nuclear isolation. **C)** Principal component analysis (PCA) of the mass spectrometry data after µMap in HEK293T cells. **D)** Of the top 40 enriched proteins in µMap, 28 (70%) are annotated interactors of c-Myc on the IntAct database. **E, F)** Gene ontology analysis of significantly enriched proteins in µMap. The molecular functions and cellular components significantly represented in the data align with known functions of c-Myc-binding co-regulatory proteins. **G)** µMap measures subtle changes to c-Myc interactions induced by treatment of JQ-1 (1µM, 37°C for 3 h). BET family proteins, BRD2/3/4, are not enriched by µMap in cells treated with JQ-1, suggesting µMap captures drug-dependent changes in c-Myc protein-protein interactions at chromatin.

#### Extended Data Figure 3 – µMap identifies cancer-specific c-Myc interactions

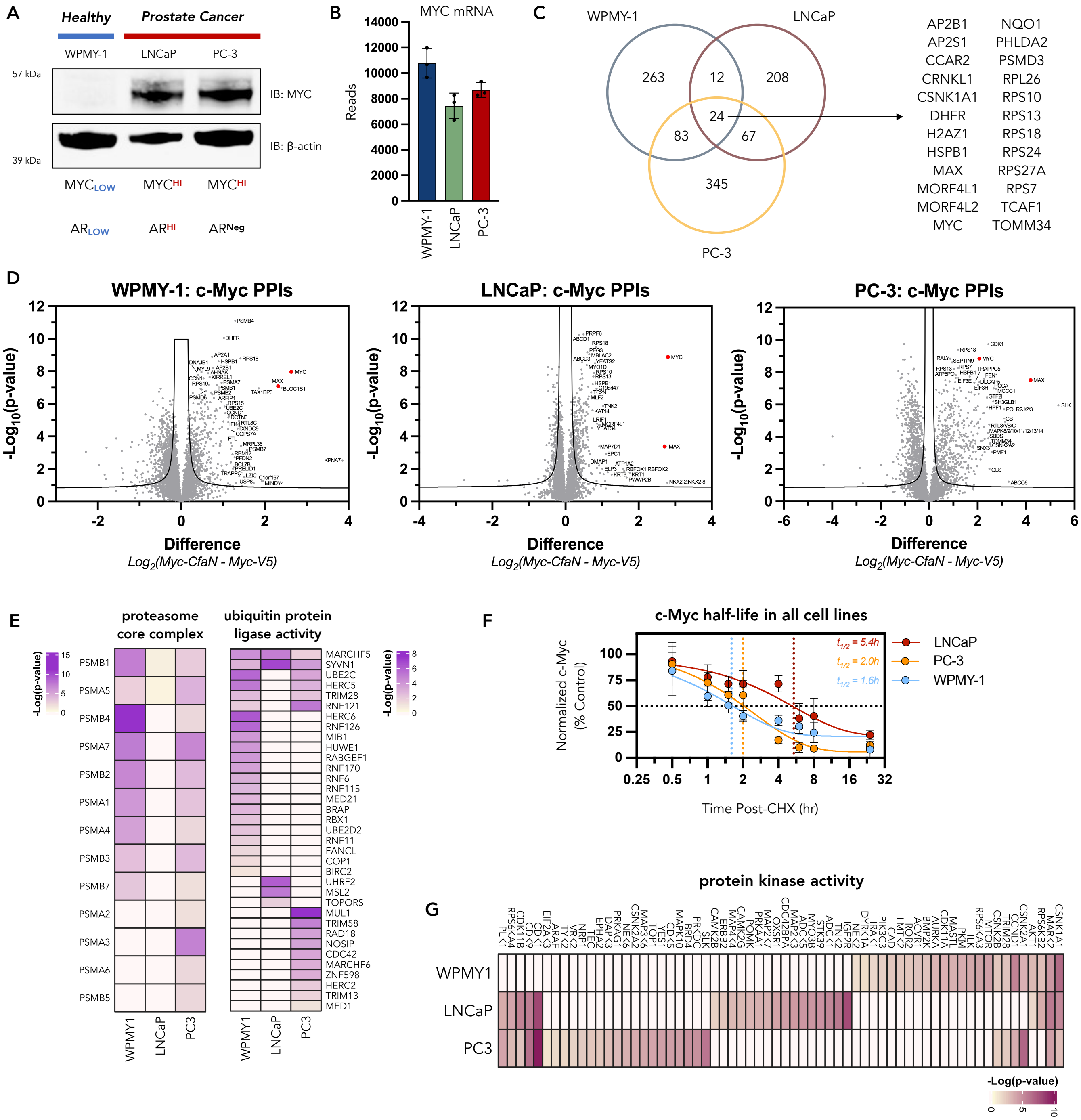


**Extended Data Figure 3. A)** Endogenous c-Myc expression in WPMY-1 (healthy), LNCaP, and PC-3 cells. **B)** Total RNA sequencing shows c-Myc mRNA is not upregulated in LNCaP or PC-3 cells, suggesting overexpression of c-Myc in cancer is not due to transcriptional upregulation of c-Myc. **C)** µMap identifies 24 common c-Myc interacting proteins across the three cell lines. **D)** Volcano plots of all proteins in mass spectrometry data after µMap in WPMY-1, LNCaP, and PC-3 cells. MYC and MAX are among the most significantly enriched proteins in each cell line. **E)** Gene ontology analyses of significantly enriched proteins related to proteasomal degradation suggest c-Myc has fewer interactions with degradation machinery in LNCaP and PC-3 cells. **F)** Cycloheximide (CHX) chase assays in each cell line suggest that c-Myc half-life is longer in LNCaP and PC-3 cells when compared to the c-Myc half-life in WPMY-1 cells. **G)** Gene ontology analyses of significantly enriched proteins with kinase activity suggest many kinases interact with c-Myc in cancer cells only.

#### Extended Data Figure 4 – SLK interacts with c-Myc and drives cancer-specific effects

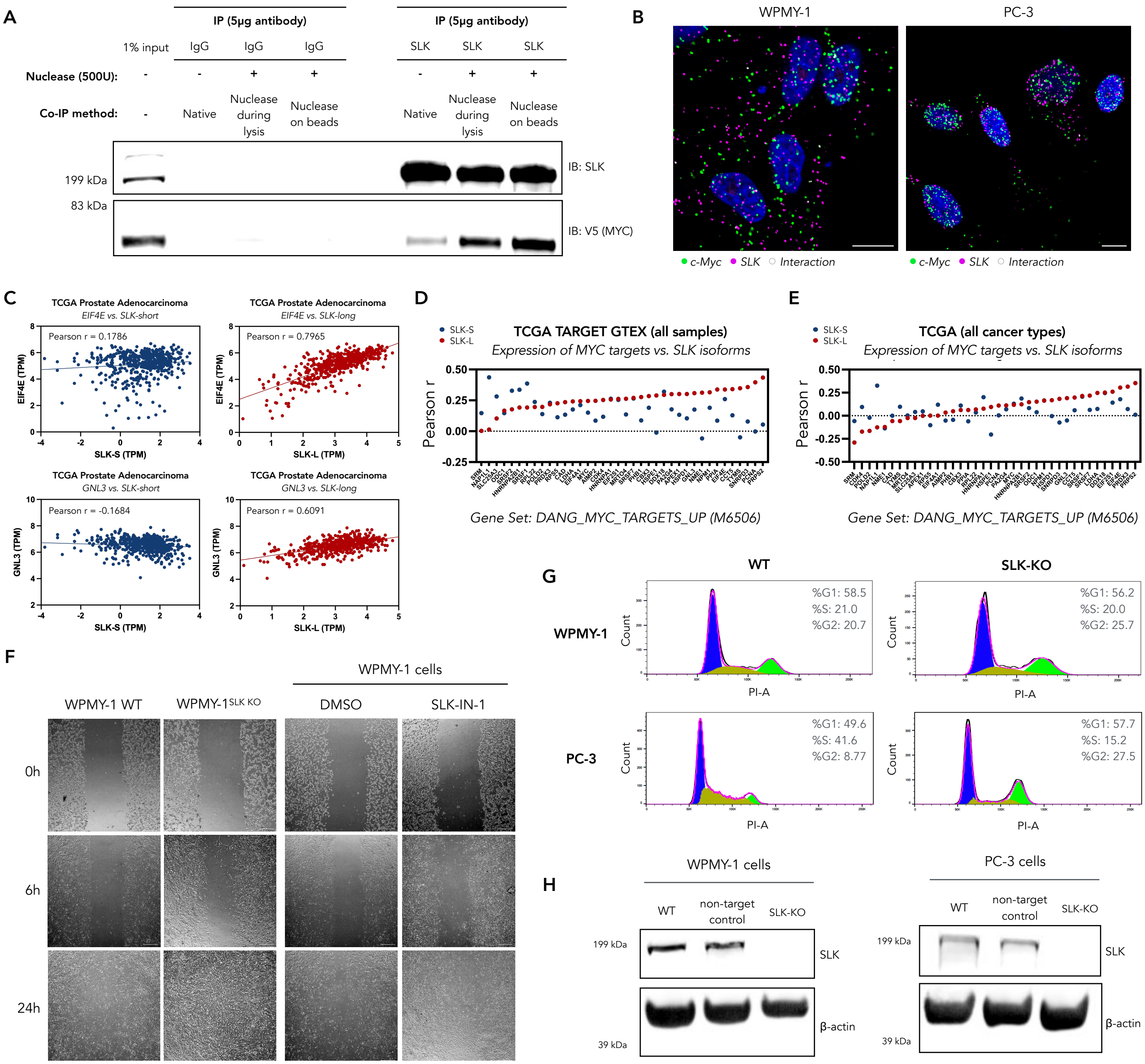


**Extended Data Figure 4. A)** Co-immunoprecipitation (co-IP) of SLK and c-Myc in PC-3 cells. Recovery of c-Myc after immunoprecipitation of SLK increases when a universal nuclease is added to the lysate, suggesting the Myc-SLK interaction is not scaffolded by DNA. **B)** Immunofluorescent staining of c-Myc (green) and SLK (magenta) with rolling circle amplification (MolBoolean™ assay) suggests c-Myc interacts with SLK in the nucleus of PC-3, but not WPMY-1 cells. Furthermore, SLK localization is primarily cytoplasmic in WPMY-1 cells, but nuclear in PC-3 cells. Scale bar = 10μm. **C-E)** Analysis of RNA sequencing data from the TCGA TARGET GTEX study on the UCSC Xena database (xenabrowser.net).[2,3] **C)** In prostate cancer samples, expression of the SLK-L isoform (red) correlates with EIF4E and GNL3 expression (Myc target genes), but this is not true for the SLK-S isoform (blue). **D)** Expression of c-Myc target genes in all samples (normal and cancer, N = 19,131) correlates with expression of the SLK-L isoform. **E)** Expression of c-Myc target genes in TCGA samples (cancer only, N = 10,535) correlates with expression of the SLK-L isoform. **F)** 24-hour scratch assays suggest that WPMY-1 cell migration is not changed by SLK knockout or a commercially available SLK inhibitor (SLK/STK10-IN-1, 10μM). Scale bar = 200μm. **G)** Cell cycle analysis with propidium iodide. WPMY-1 cell cycle is minimally affected by SLK knockout, while SLK knockout in PC-3 cells reduces the percentage of cells in S-phase and leads to G2 stalling. **H)** Western blot validation of SLK-KO WPMY-1 and PC-3 cell lines generated with CRISPR using the pSpCas9(BB)-2A-GFP (PX458) plasmid and *SLK*-specific gRNA.

### Extended Data Figure 5 – Global proteomics in WT and SLK-KO PC-3 cells

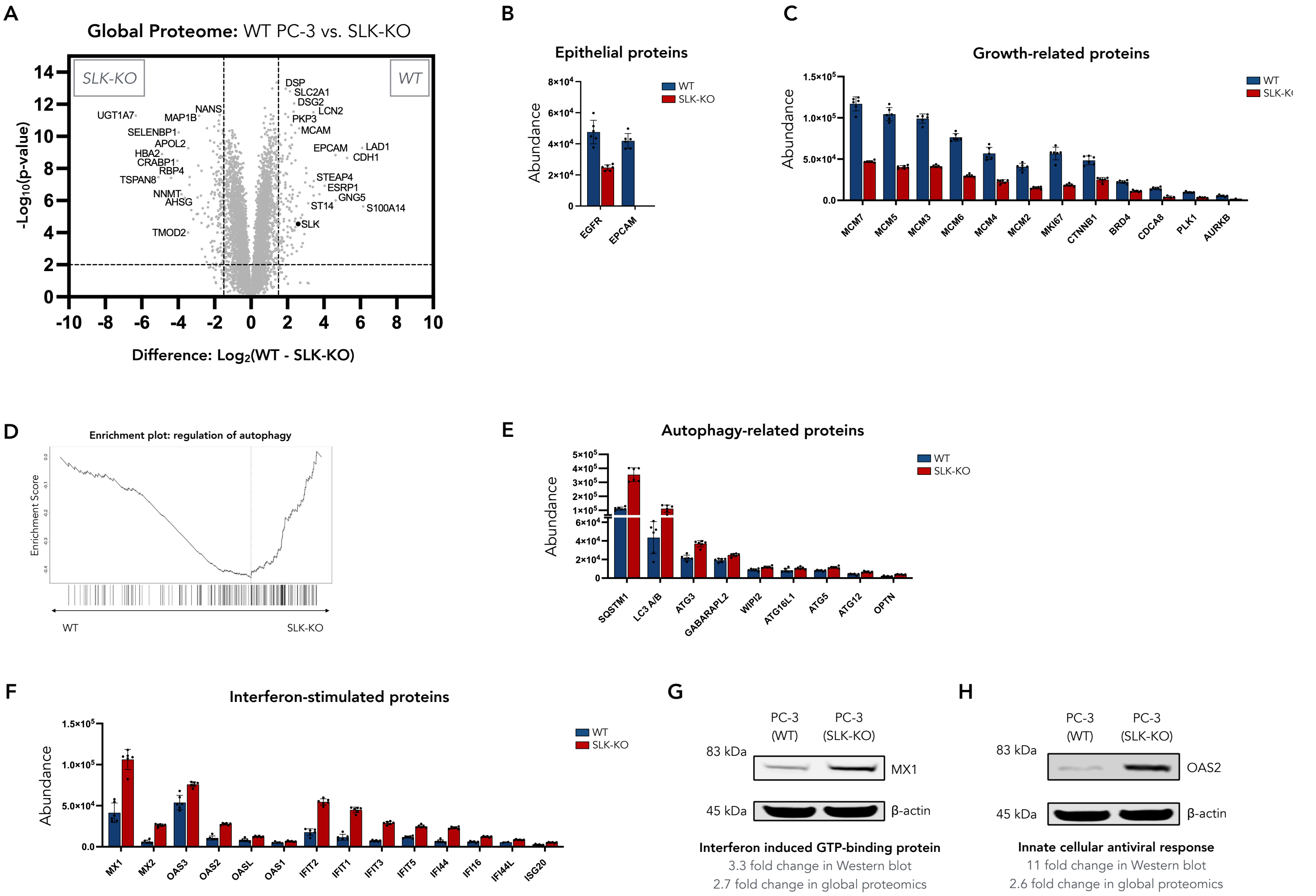


**Extended Data Figure 5.** **A)** The global proteome of WT PC-3 cells differs from the global proteome of SLK-KO PC-3 cells. **B)** WT PC-3 cells exhibit increased expression of EGFR, and EpCAM—cell surface proteins associated with epithelial cell functions. **C)** WT PC-3 cells exhibit increased expression of canonical proteins involved in DNA replication (the MCM2-7 ring of the MCM complex), proliferation (Ki-67, β-catenin, CDCA8), and the regulation of mitosis (BRD4, AURKB, PLK1). **D, E)** SLK-KO PC-3 cells exhibit increased expression of proteins that regulate autophagy. Specifically, selective autophagy receptors (p62/SQSTM1, OPTN) and complexes involved in autophagosome formation (Atg3-LC3 conjugate, Atg12-5-16L1 complex, WIPI2, GABARAPL2)[4,5] have increased expression in SLK-KO PC-3 cells. **F)** Additionally, interferon stimulated genes that are activated in response to dsDNA sensing are upregulated in SLK-KO PC-3 cells. **G, H)** Upregulation of MX1 and OAS2 was confirmed by Western blotting.

#### Extended Data Figure 6 – SLK regulates autophagy and promotes an epithelial phenotype in PC-3 cells

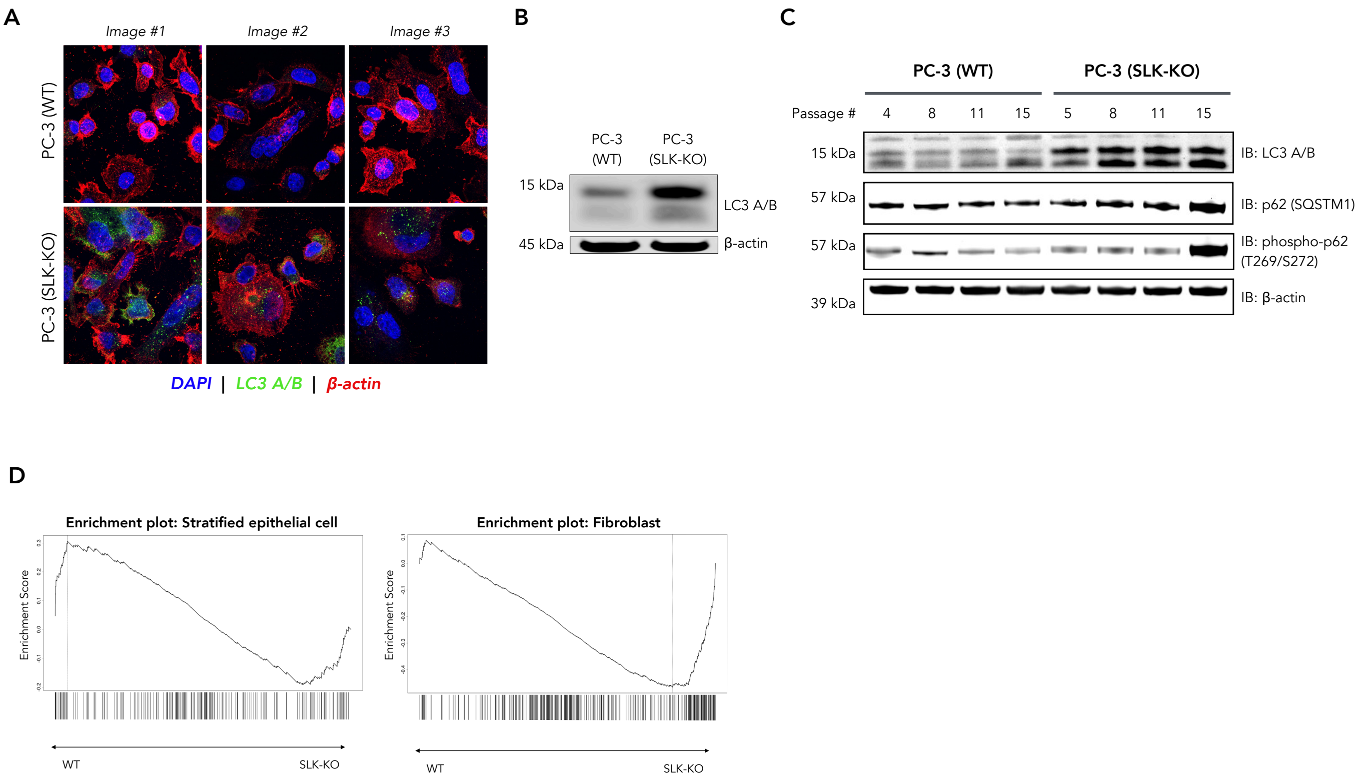


**Extended Data Figure 6. A,B)** Western blotting and immunofluorescent staining of LC3 A/B suggests LC3 protein is upregulated in SLK-KO PC-3 cells. **C)** In the SLK-KO cells, LC3 A/B protein and p62/SQSTM1 accumulate over time, but this is not observed in WT PC-3 cells. These data suggest autophagy is inhibited in SLK-KO cells, as proteins in the autophagosome are not being degraded along with the cargo, which occurs when autophagic flux is normal. **D)** Gene set enrichment analysis suggests SLK-KO induces an epithelial-to-mesenchymal transition.

#### Extended Data Figure 7 – Phosphoproteomics in WT and SLK-KO PC-3 cells

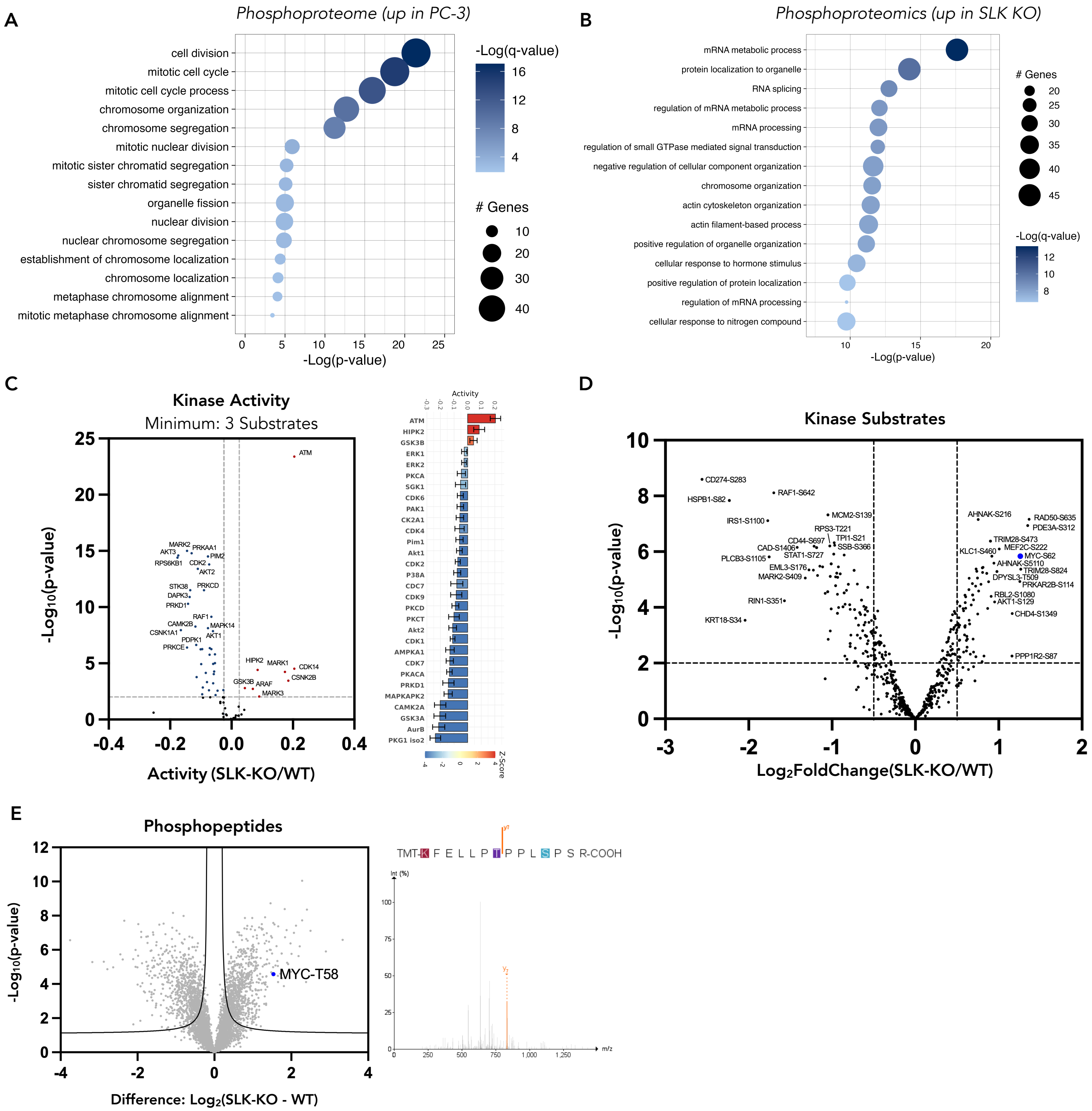


**Extended Data Figure 7. A)** Gene ontology analysis of the enriched phosphoproteome in WT PC-3 cells suggests proteins that regulate mitosis are active. **B)** Gene ontology analysis of the enriched phosphoproteome in SLK-KO PC-3 cells suggests proteins that regulate mRNA metabolism, splicing, and GTP signaling are active. **C)** The most active kinase in SLK-KO PC-3 cells is ATM, which is activated following double stranded breaks (DSBs) and regulates the DNA damage response by phosphorylating a network of substrates. Raw data was analyzed with MaxQuant, and kinase activity was predicted using RoKAI App (https://rokai.io/), using a minimum cutoff of 3 substrates. **D)** The phosphosite output from the analysis with RoKAI shows RAD50-S635—a substrate of ATM that is required for the coordination of DNA DSB repair—is enriched in the SLK-KO PC-3 cells. **E)** Additional analysis of the enriched phosphoproteomes with FragPipe shows the canonical phosphodegron of c-Myc (PT*PPLS*P) is enriched in the SLK-KO cells, suggesting SLK prevents c-Myc protein from being degraded through phosphorylation at this site. Spectra were visualized using PDV.

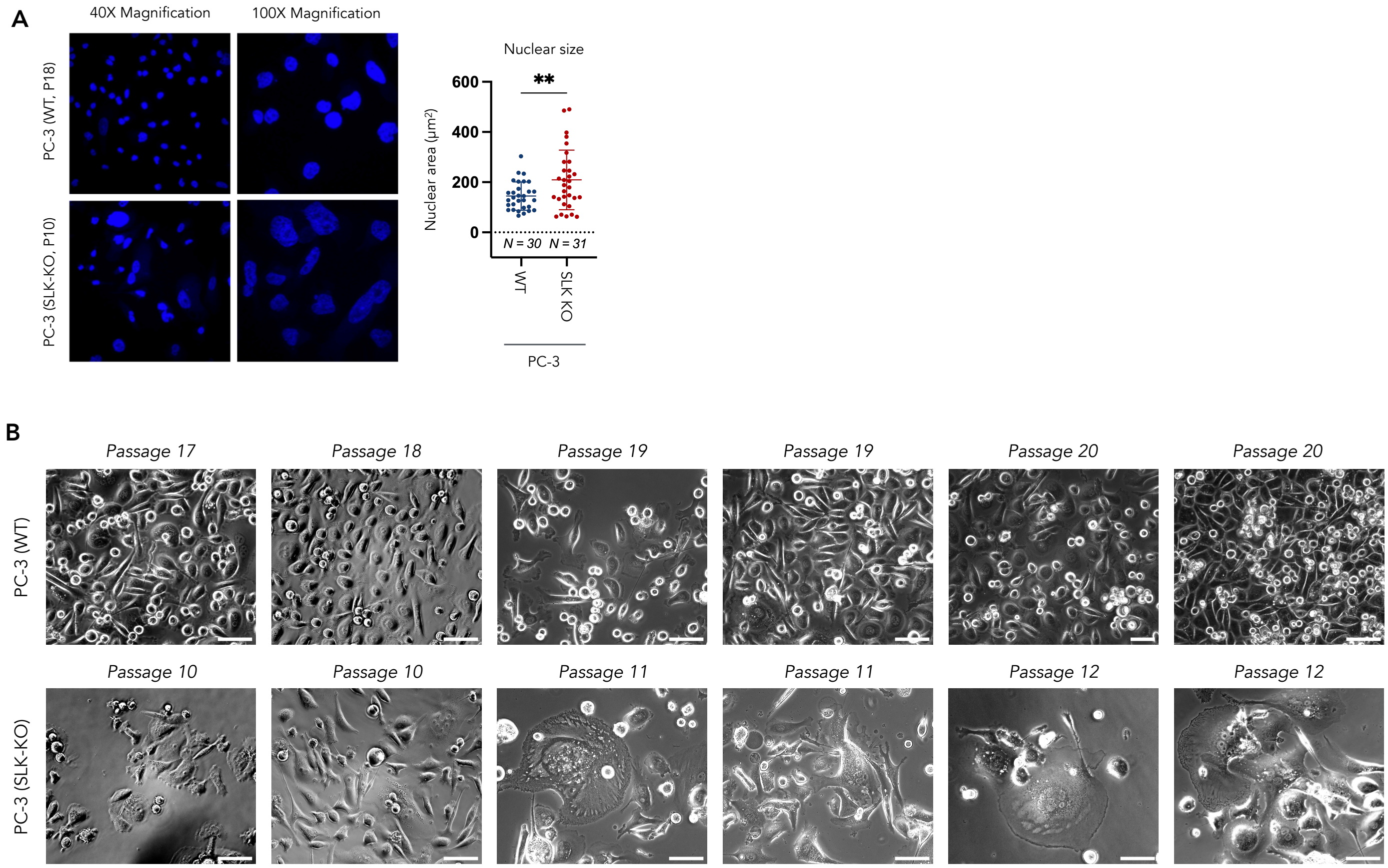


**Extended Data Figure 8. A)** The average size of the nucleus is increased in SLK-KO cells, suggesting a loss of chromatin integrity. **B)** Brightfield images of WT (top) and SLK-KO (bottom) PC-3 cells over time. WT cells appear uniform in size and morphology at all passages, but SLK-KO cells dramatically increase in size and change morphology over time, exhibiting enlarged nuclei and multinucleated cells. This phenotype suggests the SLK-KO cells may be unable to progress through metaphase due to a lack of chromatin integrity.

#### Materials and Methods

General Considerations

$Cfa^C$-Ir was synthesized as previously described.[6] The diazirine biotin probes used in μMap are commercially available (MCE, #HY-154801). TBST was purchased from Boston BioProducts (#IBB-180X). iBright Prestained Protein ladder was purchased from Thermo Scientific (#LC5615). Water was purified using a Millipore Milli-Q Integral Water Purification System.

Cell Culture

HEK293T cells were a gift from the MacMillan group (Princeton University), and other cell lines were purchased from American Type Culture Collection (ATCC). All cells were cultured in a humidified incubator at 37°C with 5% $CO_2$. HEK293T cells and WPMY-1 cells (ATCC, #CRL-2854) were maintained in DMEM (Gibco, #11995-065). LNCaP cells (ATCC, #CRL-1740) were maintained in RPMI 1640 (Corning, #10-041-CV). PC-3 (ATCC, #CRL-1435) were maintained in DMEM/F12, GlutaMAX (Gibco, #10565-018). All basal media were supplemented with 10% v/v FBS (R&D Systems, #S11150), and 100 U $mL^{-1}$ of penicillin 100 μg $mL^{-1}$ of streptomycin (Gibco, #15140-122) to make the complete growth medium in which cells were cultured.

Western blot

Cell lysates were quantified using Pierce BCA protein assay kit (Thermo Scientific, #A55864), normalized to the same concentration, and 4X Laemmli Sample Buffer (BioRad, #1610747) containing 2-mercaptoethanol was added to reach a 1X final concentration. Samples were run on a pre-cast Invitrogen NuPAGE 4-12% bis-tris acrylamide gel (Invitrogen #NW04125BOX) and transferred to nitrocellulose membrane (Thermo Scientific, #88018). Membranes were blocked with 5% w/v milk in TBS-T (25 mM tris, 150 mM NaCl, 0.1% v/v Tween-20, pH 7.7) for 1 h at room temperature. Membranes were incubated with primary antibodies at the stated dilutions (see Table 2) overnight with rotation at 4°C. After 3x 5 min washes with TBS-T, secondary antibodies were applied for one hour at room temperature (see Table 2), before imaging on a Li-Cor Odyssey imager (LI-COR; IRDye secondaries) or an ImageQuant LAS 500 (GE Healthcare; HRP secondaries). Quantification of Western blots was performed in Fiji.

Immunofluorescence

Cells were harvested with trypsin, washed with PBS, plated on polylysine-treated coverslips, then incubated overnight at 37°C. Cells were then fixed with 4% paraformaldehyde diluted in PBS for 10 min at room temperature. After washing 3x with PBS, cells were permeabilized with 75% ethanol diluted in PBS overnight at 4°C. The coverslips were then blocked with 1% BSA in PBS for 1 h at room temperature. Coverslips were then incubated with primary antibody at the stated dilutions (see Table 2) in 1% BSA in PBS overnight at 4°C. Samples were then washed 3x with PBS and incubated with secondary antibody diluted 1:1000 for 1 h at room temperature. Slides were then washed 3x with PBS and mounted to slides using ProLong™ Gold Antifade Mountant with DNA Stain DAPI (Invitrogen, #P36935). Images were acquired on the FLUOVIEW FV3000 confocal microscope (Olympus).

Total RNA sequencing

WPMY-1, LNCaP, and PC-3 cells were seeded in 6-well plates to reach 60-75% confluency the following day, then incubated overnight at 37°C with 5% $CO_2$. For the transfected PC-3 cell conditions, the media was replaced with fresh media, and PC-3 cells were transfected with either c-Myc-CfaN (0.75μg) or c-Myc-V5-6xHis (0.75μg) using the Lipofectamine 3000 Transfection Reagent (Thermo Scientific, #L3000150), following the manufacturer's instructions. The next day, cells were harvested with trypsin, washed with PBS, and RNAs were extracted from $1x10^6$ cells using the Rneasy plus mini kit (Qiagen, #74134) according to the manufacturer's instructions. Total RNA was submitted to The Herbert Wertheim UF Scripps Institute for Biomedical Innovation & Technology - Genomics Core (RRID:SCR_017827), where it was quantified using a Qubit 2.0 Fluorometer (Invitrogen, Carlsbad, CA) and evaluated on an Agilent 4200 TapeStation (Agilent Technologies, Santa Clara,

CA) for quality assessment. All RNA samples had RNA Integrity Number (RIN) = 10.0 and were used for total RNA-seq library preparation.

RNase-free working environment was maintained, and RNase-free tips, tubes, and plates were utilized. 1µg of total RNA per sample was depleted of ribosomal RNA using probes provided in the NEBNext rRNA Depletion Kit v2 (Cat. #E7405, NEB, Ipswich, MA) according to the manufacturer's recommendations. The library preparation from the rRNA-depleted RNA was conducted according to the NEBNext Ultra II Directional RNA kit (Cat. #E7760, NEB, Ipswich, MA). Briefly, the RNA samples were chemically fragmented in a buffer containing divalent cations by heating to 94°C for 10 min. The fragmented RNA was random hexamer primed and reverse transcribed to generate the first strand of cDNA. The second strand was synthesized after removing the RNA template and incorporating dUTP in place of dTTP to maintain strand specificity. Fragmented ds cDNA was then end repaired and adenylated at their 3' ends. A corresponding 'T' nucleotide on the adaptors was utilized for ligating the adaptor sequences to the cDNA. The adaptor ligated DNA was purified using magnetic beads and PCR amplified (using 11 cycles) to incorporate a unique barcode and to generate final libraries. The libraries were purified using magnetic beads to remove any remaining primers and adaptors.

The final libraries were validated on an Agilent 4200 TapeStation (Agilent Technologies, Santa Clara, CA), normalized to 4nM, pooled equally, and loaded onto an Illumina NextSeq 2000 P3 300-cycle flow cell (Cat. # 20040561, Illumina, San Diego, CA) at 750pM final concentration and sequenced using 2 x 155bp paired-end chemistry. On average, we generated 86 million reads pass filter per sample. Raw and processed data files were uploaded to the NCBI Gene Expression Omnibus (GSE307813).

Analysis of RNA sequencing data

1. *Splicing analysis*

RNA sequencing data were processed using the nf-core/rnasplice pipeline (version 1.0.4, doi: 10.5281/zenodo.8424632) on the HiPerGator high-performance computing cluster with Nextflow (version 25.04.4) and Singularity (version 3.10.4). Raw FASTQ files were quality-checked with FastQC (version 0.12.1) and trimmed using TrimGalore (version 0.6.7, cutadapt version 3.4). Reads were aligned to the ENSEMBL human genome GRCh38 release 111 (Homo_sapiens.GRCh38.dna_sm.primary_assembly.fa) using STAR (version 2.7.9a) with the corresponding STAR index. Gene annotations were derived from the GTF file (Homo_sapiens.GRCh38.111.gtf). FeatureCounts (subread version 2.0.1) was used to quantify read counts, followed by differential expression analysis with edgeR (R version 4.0.3). Event-based differential splicing analysis was performed using rMATS (version 4.1.2).

2. *Analysis of Myc target gene expression after transfection of PC-3 cells*

Version 3.12.0 of the nf-core/rnaseq pipeline (doi: 10.5281/zenodo.1400710) was run on HiPerGator with Nextflow (v25.04.4) and Singularity (v3.10.4). All default settings were used except the following changes: the option for alignment by STAR was used, along with RSEM for quantification, and reads were aligned to the human GRCh38 genome. Downstream analysis was performed with DESeq2 (v1.34.0) to identify differentially expressed (DE) genes.

Analysis of publicly available RNA sequencing data

The analyses of publicly available RNA sequencing data in this manuscript are based upon data generated by the TCGA Research Network, the Therapeutically Applicable Research to Generate Effective Treatments (TARGET) initiative, and The Genotype-Tissue Expression (GTEx) Project (see Acknowledgements). The data from the TCGA TARGET GTEx study were obtained from the UCSC Toil RNAseq Recompute Compendium,[2,3] and further analyzed and plotted in GraphPad Prism 10. The human gene set: DANG_MYC_TARGETS_UP was used to select Myc target genes in the analysis.[7]

µMap photoproximity labeling

All µMap experiments were performed with three biological replicates per group (with 2-4x 15cm plates per replicate).

1. *Transfection*

Cells were seeded in 15cm plates to reach 60-75% confluency the following day, then incubated overnight at 37°C with 5% $CO_2$. The media was replaced with fresh media, and all plates were transfected with either c-Myc-V5-6xHis (control) or c-Myc-CfaN (µMap) using the Lipofectamine 3000 Transfection Reagent (Thermo Scientific, #L3000150) and following the manufacturer's instructions. 18-24 h post-transfection, the cells were harvested using trypsin, washed once with PBS, and pelleted in 15mL conical vials before snap-freezing on dry ice and storage at -80°C.

2. *µMap photoproximity labeling*

The next day, cell pellets were thawed and resuspended in 0.2mL Nuclear Isolation Buffer (NIB) per $1x10^6$ cells (NIB: 45mM KCl, 0.1mM EDTA, 6.25mM $MgCl_2$, 12.5mM Tris-HCl (pH 7.5), 375mM sucrose, 0.125% NP-40, and 1X halt protease inhibitor cocktail). Cell suspensions were incubated on ice for exactly 7 min, then centrifuged at 600xg for 10 min at 4°C to isolate crude nuclei. The supernatant was discarded, and nuclei were washed twice with PBS, centrifuging at 400xg for 3 min at 4°C. The nuclei were then treated with 1.0µM CfaC-Ir diluted in PBS and incubated in a 37°C water bath for 40 min with gentle shaking every 10 min while protected from light with foil. The nuclei were pelleted by centrifugation at 400xg for 3 min at 4°C, then washed 3x with PBS. Nuclei were resuspended in 1mL of 250µM diazirine-PEG3-biotin conjugate and irradiated for 1 min at 450nm with 100% light intensity on with the Photoreactor m2 (Acceled Bio). Nuclear pellets were then washed 2x with PBS, centrifuging at 400xg for 3 min. Nuclear pellets were lysed in LB3 buffer (1mM EDTA, 0.5mM EGTA, 10mM Tris-HCl (pH 7.5), 100mM NaCl, 0.1% Na-Deoxycholate, 0.5% N-lauroyl sarcosine, 1X HALT protease inhibitor cocktail) and sonicated using the Diagenode Bioruptor Sonicator (HIGH, 14 cycles at 4°C, 30 s ON, 30 s OFF). Lysates were then centrifuged at 17,000xg for 20 min at 4°C. The supernatant was extracted, and protein concentrations were determined using the BCA assay and normalized to 1 mg/mL in 1mL total volume using LB3 buffer. Lysates were incubated overnight at 4°C with 100µL pre-washed Streptavidin Mag Sepharose beads (Cytiva, #28985799) per 1mg lysate with end-over-end rotation.

3. *Washing*

Beads were washed 3x 5 min with 1% SDS in PBS (rotating), then 3x 5 min with 1M NaCl in PBS (rotating), followed by 3x 5 min with 10% EtOH in PBS (rotating). The beads were then transferred to new Lo-bind tubes, and washed 3x with PBS and 3x with 100mM ammonium bicarbonate. The beads were then resuspended in 0.5mL 3M urea in PBS, and 25µL dithiothreitol (200mM in 25mM ammonium bicarbonate) was added to each sample to reduce proteins. Samples were incubated at 55°C for 30 min with shaking at 700rpm. Next, 30µL iodoacetamide (500mM in 25mM ammonium bicarbonate) was added to each sample and incubated at 30 min at room temperature in the dark to alkylate proteins. Beads were then washed 3x with PBS and 6x with 50mM ammonium bicarbonate, then transferred to new Lo-bind tubes. To elute peptides, the beads were resuspended in 40µL of 50mM ammonium bicarbonate with 1.2µL MS-grade trypsin (resuspended in 1mg/mL in 50mM acetic acid). Samples were incubated overnight at 37°C with end-over-end rotation, followed by another addition of 0.8µL trypsin for 1 hour at 37°C the next morning. Supernatants were collected and split into technical replicates of 20µL each before storage at -80°C.

Label free proteomics

Peptide digests were acidified with TFA to 0.1% (v:v) and desalted using 2 µg capacity ZipTips (Millipore, Billerica, MA) according to manufacturer instructions. Following drying under vacuum, peptides were re-solubilized in 0.1% formic acid (FA) to a final concentration of 100ng/µL. Samples were analyzed on a nanoElute (plug-in V2.1.60.0; Bruker, Bremen, Germany) coupled to a Bruker TimsTOF Pro 2 mass spectrometer (Breman, Germany), equipped with a CaptiveSpray source and a 20µm zero dead volume (ZDV) Sprayer. Peptides (corresponding to 100ng) were loaded onto a Thermo Fisher (Waltham, MA) PepMap Neo C18 trap column (300mm X 5mm, 5mm particle size) and then separated on a PepSep Series reverse-phase C18 column (15cm X 150µm, 1.5µm particle size) from Bruker (Bremen, Germany). The column temperature was maintained at 50°C using an integrated Bruker Column Toaster (Bremen, Germany). The column was equilibrated using 4 column volumes before loading samples in 100% buffer A (99.9% Fisher Optima® LC/MS water, 0.1% FA), with both steps performed at 800 bar. The trap column was equilibrated at 201.6 bar. Samples were separated at 500nl/min using a linear gradient from 2% to 35% buffer B (99.9% Fisher Optima® LC/MS acetonitrile, 0.1% FA) over 20.0 min before ramping to 95% buffer B (in 0.5 min) and sustained at 95% buffer B for 4.5 min (total separation method time 25.0 min). The Bruker TimsTOF Pro 2 was operated in DIA-PASEF mode using Tims Control v. 5.0.2. Settings for the MS method were as follows: Mass Range 100 to 1700m/z, 1/K0 Start 0.6 V.·/cm2 End 1.4 V·s/cm2, TIMS Ramp and accumulation time 75ms, Capillary Voltage 1700V, Dry Gas 3 l/min, Dry Temp 200°C, DIA-PASEF settings: 18 MS/MS scans (50m/z windows, 0.21 1/K0 windows, total cycle time 0.74), mass range 300 to 1200, and CID collision energy 20eV (at 0.60, 1/K0) to 65eV (at 1.60, 1/K0). The analysis was performed at The Herbert Wertheim UF Scripps Institute for Biomedical Innovation & Technology, Mass Spectrometry and Proteomics Core Facility (RRID:SCR_023576).

Label free proteomics data analysis

The raw mass data were further analyzed by DIA-NN (1.8.1). Parameters set as follows: trypsin/P digestion, missed cleavages: 3, maximum number of variable modifications: 3, N-term M excision, Ox(M), Ac(N-term), and C carbamidomethylation. Peptide length range was 7-30, precursor charge range 1-4, m/z range 300-1800, and fragment ion range 200-1800. Mass accuracy and MS accuracy were both set to 10. The following settings on the algorithm were checked: "Use isotopologues", "MBR", "No shared spectra", and "Heuristic protein inference". Precursor FDR was set to 1% with threads as 16. A spectral library was generated via DIANN from all known human proteins (In-Silico spectral library). The resulting matrix.pg file was opened in Perseus (v2.0.7.0). Intensities were inputted as "main", and the rest of the descriptors are categorical. Data was then transformed (Log2). Data is annotated by treatment. Missing values were imputed with Perseus default settings (0.3 Width, 1.8 Down shift). Normalization was performed via median subtraction. Following this process, a volcano plot was generated utilizing a t-test for statistical significance. The resulting volcano plots were plotted in GraphPad Prism 10 for final figures.

Global proteomics

Wild-type PC-3 cells and SLK-KO PC-3 cells were seeded in 10cm plates to reach 80-100% confluency the following day, then incubated overnight at 37°C with 5% $CO_2$. Each cell line was plated in three 10cm plates, for three biological replicates per group. The next morning, the cells were harvested using trypsin, washed once with PBS, and pelleted in 15mL conical vials before lysing with SDS lysis buffer (50mM HEPES, pH 8.5, 1% SDS, 1X halt protease inhibitor cocktail). 20 µg of protein was taken for cleanup and digestion. Proteins were cleaned up and digested using Sera-Mag Carboxylate SpeedBeads (Cytiva Life Sciences: E7 Cat. #45152105050250 and E3 Cat. #65152105050250) following the manufacturer's protocol. Digested peptides were split into two technical replicates per sample. Label free proteomics and subsequent data analysis were performed as described above.

Phospho proteomics

1. *Transfection of c-Myc-V5-6xHis*

Wild-type PC-3 cells and SLK-KO PC-3 cells were seeded in 10cm plates to reach 60-75% confluency the following day, then incubated overnight at 37°C with 5% $CO_2$. Each cell line was plated in five 10cm plates, for five biological replicates per group. The next morning, the media was replaced with fresh media, and all plates were transfected with c-Myc-V5-6xHis (1µg) using the Lipofectamine 3000 Transfection Reagent (Thermo Scientific, #L3000150), following the manufacturer's instructions.

2. *Cell lysis, protein precipitation, and digestion*

Phosphopetides were enriched as previously described.[8] Briefly, 18-24 h post-transfection, the cells were harvested using trypsin, washed once with PBS, and pelleted in 15mL conical vials before lysing with lysis buffer (200mM HEPES, pH 8.5 and 8M urea with 1X Halt Protease and Phosphatase Inhibitor Single-Use Cocktail (Thermo Scientific, #1861280)). The lysates were then passed through a 21-gauge needle 10 times. Protein concentrations were determined by the Pierce BCA protein assay kit (Thermo Scientific, #A55864), and all samples were normalized to 1 mg/mL. Disulfide bond reduction was performed by adding 5mM tris (2-carboxyethyl) phosphine (TCEP) to each sample and incubating for 15 min at room temperature. Alkylation was performed by then adding 10mM iodoacetamide (IAA) to the samples and incubating for 30 min in the dark. Excess IAA was quenched by adding 10mM dithiothreitol (DTT) to the samples and incubating for 15 min at room temperature. 400µL of 100% methanol was added to 100µg of each sample (0.1mL) in a 1.5mL microcentrifuge tube, and the samples were vortexes for 5 s. 100µL of 100% chloroform was added to each sample, and the sample was vortexed for 5 s. 300µL of water was added to each sample, and the sample was vortexed for 5 s. Samples were centrifuged for 1 min at 14,000xg. The aqueous and organic phases were removed, isolating the protein disk, which was then washed with 400µL of 100% methanol and centrifuged at 21,000xg for 2 min at room temperature. The supernatant was removed and samples were resuspended in 70µL 200mM HEPES, pH 8.5. The samples were digested overnight at 37°C with trypsin (1µg per sample, Thermo Scientific #90057).

3. *TMT labeling*

30µL acetonitrile was added to the digested peptides, and the samples were vortexed. TMT10plex Mass Tags (Thermo Scientific, #90111) were prepared according to the manufacturer's instructions; samples were labeled with 200µg of TMT reagent, which was incubated for 1 h at room temperature with the peptides. The mass tags added to each sample were recorded. The reaction was quenched by adding 0.66µL of 50% hydroxylamine in water (final concentration of 0.3% hydroxylamine v/v). The TMT-labeled samples were pooled together at a 1:1 ratio, and the volume was evaporated down to ~30% of the total volume by vacuum centrifuge. The sample was then desalted with Pierce Peptide Desalting Spin Columns (Thermo Scientific, #89852) according to the manufacturer's instructions.

4. *Phosphopeptide enrichment*

Phosphopeptides were enriched using the Pierce High-Select Fe-NTA Phosphopeptide Enrichment Kit (Thermo Scientific, #A32992) according to the manufacturer's instructions.

5. *TMT quantitative proteomics and mass spectrometry*

Peptide digest samples corresponding to the enriched phosphoproteome, and the flow-through proteome following the enrichment were dried under vacuum and subsequently acidified using 100 mL of 1% TFA (pH < 3). The samples were then desalted using NuTip Carbon tips from Glygen (Glygen Corp., Columbia, MD) in the case of phosphopeptides, or 2µg capacity ZipTips (Millipore, Billerica, MA), according to manufacturer instructions, respectively. They were then again dried under vacuum. Peptides resolubilized in 5mL of 0.1% TFA were on-line eluted into a Fusion Tribrid mass spectrometer (Thermo Scientific, San Jose, CA) from an EASY PepMap™ RSLC C18 column (2µm, 100Å, 75µm x 50cm, Thermo Scientific, San Jose, CA), using a gradient of 5-25% solvent B (80/20 acetonitrile/water, 0.1% formic acid) in 180 min, followed by 25-44% solvent B in 60 min, 44-80% solvent B in 0.1 min, a 5 min hold of 80% solvent B, a return to 5% solvent B in 0.1 min, and finally a 10 min hold of 5% solvent B. The gradient was then extended for the purpose of cleaning the column by increasing solvent B to 100% in 3 min, a 100% solvent B hold for 10 min, a return to 5% solvent B in 3 min, a 5% solvent B fold for 3 min, an increase of solvent B to 100% in 3 min, a 100% solvent B hold for 10 min, a return to 5% solvent B in 3 min and a 5% solvent B hold for 3 min and finally, another increase to 100% solvent B in 3 min and a hold of 100% solvent B for 10 min. All flow rates were 250nL/minute delivered using a Thermo Vanquish Neo UHPLC nano liquid chromatography system (Thermo Scientific, San Jose, CA). Solvent A consisted of water and 0.1% formic acid. Ions were created at 2.1kV using an EASY Spray source (Thermo Scientific, San Jose, CA) held at 50 °C. A synchronous precursor selection (SPS)-MS3 mass spectrometry method was selected based on the work of Ting et al.[9] Scans were conducted between 380-2000 m/z at a resolution of 120,000 for MS1 in the Orbitrap mass analyzer at an AGC target of 4E5 and a maximum injection of 50 msec. Collision induced dissociation (CID) was then performed in the linear ion trap of peptide monoisotopic ions with charge 2-8 above an intensity threshold of 5E3, using a quadrupole isolation of 0.7 m/z and a CID energy of 35%. The ion trap AGC target was set to 1.0E4 with a maximum injection time of 50 msec. Dynamic exclusion duration was set at 60 s and ions were excluded after one time within the +/- 10ppm mass tolerance window. The top 10 MS2 ions in the ion trap between 400-1200 m/z were then chosen for Higher-energy C-trap dissociation (HCD) at 65% energy. Detection occurred in the Orbitrap at a resolution of 60,000 and an AGC target of 1E5 and an injection time of 120msec (MS3). All scan events occurred within a 3-second specified cycle time. The analysis was performed at The Herbert Wertheim UF Scripps Institute for Biomedical Innovation & Technology, Mass Spectrometry and Proteomics Core Facility (RRID:SCR_023576).

6. *Data analysis with FragPipe*

Raw data were converted to mzmL files using MSConvert, following the online tutorial (https://fragpipe.nesvilab.org/docs/tutorial_convert.html). Converted data were then processed using FragPipe using the TMT10-MS3 workflow with indicated 229.16293 modification for TMT label adjusted for N-terminus. Samples were filtered by "Reverse" and "Potential contaminant" and were categorized by treatment. A FASTA file of uniprot Homo sapiens proteome was included as the reference. The protein abundance file was processed through Perseus, and protein were filtered by PSM>1. Following this process, a volcano plot was generated utilizing a t-test for statistical significance. The resulting volcano plots were plotted and analyzed following the procedures listed in the label free proteomic workflow.

7. *Data analysis with MaxQuant*

The analysis with MaxQuant was executed as previously described.[10] The Phospho(STY) sites file was loaded into Perseus. Rows were filtered based on "Reverse", "Potential contaminant", and localization probability > 0.75. Data was then transformed ($Log_2$). The data table was lengthened using "Expand site table". Samples were annotated by treatment using categorical annotation. Rows were filtered by rows containing > 50% valid values in each group. Imputation was performed using Perseus default settings. Samples were normalized by median subtraction. A volcano plot was generated utilizing a t-test for statistical significance. The resulting volcano plots were plotted in GraphPad Prism 10 for final figures. Kinase activity was predicted using RoKAI App (https://rokai.io/). Spectra were visualized using PDV (https://github.com/wenbostar/PDV?tab=readme-ov-file).

Cycloheximide chase assay

Cells were seeded in 6-well plates to reach 80-100% confluency the following day, then incubated overnight at 37°C with 5% $CO_2$. The media was then removed and replaced with media containing cycloheximide (50µg/mL). At post-treatment collection time points (0, 15, 30, 60, 90, 120, 240, 360 min), cells were washed with 1mL PBS, then lysed directly in the well with ice cold lysis buffer [M-PER (Thermo Scientific, #78501), 1X protease inhibitor (Thermo Scientific #1861279), 500U Pierce Universal Nuclease for Cell Lysis (Thermo Scientific, #88702)]. Western blotting was then performed to quantify the amount of the target protein remaining at each time point.

Native co-immunoprecipitation (co-IP)

Cells were grown in 15cm plates until 80-100% confluent. The cells were washed 2-3x with ice-cold PBS, then lysed directly in the plate with EBC buffer (50mM Tris-HCl, 120mM NaCl, 0.1mM EDTA, 0.5% NP-40, 10% glycerol, 1X protease inhibitor cocktail) and harvested via scraping. After lysing for 30 min on ice, the lysates were sonicated on the Diagenode Bioruptor Sonicator (LOW, 5 cycles at 4°C, 7 s ON, 5 s OFF). Samples were then centrifuged (12,000xg, 15 min, 4°C), the supernatant was harvested, and all lysates were normalized to 1 mg/mL. 1-5 µg of the primary antibody and the isotype control antibody were then conjugated to Pierce Protein A Magnetic Beads (Thermo Scientific, #88845) according to the manufacturer's instructions. Then lysates were then incubated with the prepared protein A beads overnight at 4°C with rotation. Beads were washed 4x with Wash buffer (50mM Tris-HCl, 150mM NaCl, 5mM EDTA, 0.1% Tween-20) and transferred to new tubes. Proteins were eluted by boiling beads at 100°C for 5 min in 1X Laemmli sample buffer diluted in water. Target proteins were detected by Western blot, using a Clean-Blot IP Detection Reagent (Thermo Scientific, #21230) to limit background associated with heavy and light chains of the antibodies. For Co-IP experiments with nuclease treatment, 500U Pierce Universal Nuclease for Cell Lysis (Thermo Scientific, #88702) was added to the lysate.

RT-qPCR

$1x10^6$ cells were harvested with trypsin, washed with PBS, and RNAs were extracted using TRIzol reagent (Invitrogen, #15596026) according to the manufacturer's instructions. The RNA samples were treated with DNase (Invitrogen, #AM1907) to remove genomic DNA. The RNA concentrations were measured using Qubit RNA BR Assay Kit (Thermo Scientific, #Q10210) and normalized to the same concentration. cDNAs were synthesized with The SuperScript III First-Strand Synthesis System (Invitrogen, #18080051) according to the manufacturer's protocol with random hexamers. The cDNA concentration was measured with Qubit ssDNA Assay Kit (Thermo Scientific, #Q10212) and normalized to 5ng/µL. 10ng of cDNA was loaded in real-time qPCR reactions, which were performed in triplicate using the Power SYBR™ Green PCR Master Mix (Thermo Scientific # 4367659) and specified primer pairs (see Table 1). Quantification of gene expression was measured using the 5 Real-Time PCR System.

CRISPR knockout cell lines

1. *Cloning into pSpCas9(BB)-2A-GFP*

The design of guide sequences targeting human *SLK* (see Table 1) was done using the CRISPick web tool of the GPP at the Broad Institute. Guide RNA sequences were cloned into pSpCas9(BB)-2A-GFP (PX458) (Addgene, #48138). The PX458 vector was digested with BbsI overnight at 37°C, followed by gel purification of the digested plasmid. 100µM of forward and reverse guide oligos were phosphorylated with T4 polynucleotide kinase (NEB, #M0201S) and annealed in a thermocycler (30 min 37°C, 5 min 95°C, ramp down 4°C at 4°C/min). The annealed guide oligos were diluted 20X in nuclease-free water and ligated into the BbsI-digested PX458 vector at 16°C overnight. NEB 5-alpha Competent E. coli (NEB, #C2987H) were transformed with the ligated product to obtain the PX458 plasmids containing guide RNA sequences targeting human *SLK* (PX458-SLK).

2. *Transfection of target cells*

Cells were seeded in 10cm plates to reach 60-75% confluency the following day, then incubated overnight at 37°C with 5% $CO_2$. The media was replaced with fresh media, and all plates were transfected with using the Lipofectamine 3000 Transfection Reagent (Thermo Scientific, #L3000150) and following the manufacturer's instructions. 18-24 h post-transfection, the cells were harvested using trypsin, washed once with PBS, and resuspended in FACS buffer (PBS with 1% BSA).

3. *Single-cell sorting and validation of knockout clones*

GFP positive cells were sorted using the BD FACSAria™ III instrument, and one cell was seeded into each well of a 96-well plate containing complete growth media. The clones were incubated at 37°C with 5% $CO_2$ until the clonal populations expanded enough to be re-plated in 10cm plates. Knockout of the target gene was then validated by Western blotting.

MolBoolean assay

The MolBoolean assay (Atlas Antibodies, #MolB00001) was executed according to the manufacturer's instructions with a mouse monoclonal antibody targeting c-Myc and a rabbit polyclonal antibody targeting SLK (see Table 2). Briefly, cells were harvested with trypsin, washed with PBS, plated on polylysine-treated coverslips, then incubated overnight at 37°C. Cells were then fixed with 3.7% paraformaldehyde diluted in PBS for 15 min at room temperature. After washing 3x with PBS, cells were permeabilized with 0.2% Triton X-100 in 1X TBS for 15 min at room temperature. The cells were washed with TBST 2x for 2 min at room temperature with rocking. The cells were incubated in blocking solution for 1 h at room temperature, rinsed with TBST, and primary antibodies were incubated on the cells overnight at 4°C. Cells are washed with TBST 3x for 3 min at room temperature, and Proximity probes A and B are diluted 1:80 with Complete Diluent and incubated on the cells for 30 min at 37°C. After washing with HSBT 1x for 3 min, and TBST 2x for 3 min, Circle oligos are hybridized to the Proximity probes by incubating the cells with Circle oligos diluted in 1X Buffer A with 1X Additive for 30 min at 37°C. The cells were washed with TBST 1x for 3 min at room temperature, then the cells were incubated with 1X Nickase enzyme diluted in 1X Buffer B for 1h at 37°C. The cells were washed with TBST 3x for 3 min at room temperature, then Tag oligos and Additive were diluted in TBS to 1X concentration and incubated on the cells for 30 min at 37°C. The Tag oligo solution was decanted off and, without washing, 1X Ligase enzyme with 1X Additive diluted in 1X Buffer A was incubated on the cells for 30 min at 37°C. After washing with HSBT 1x for 3 min, and TBST 1x for 3 min, Rolling Circle Amplification (RCA) was initiated by adding 1X Polymerase enzyme diluted in 1X Buffer C to the cells and incubating for 90 min at 37°C. The cells were then washed with TBST 2x for 5 min and 1X Detection oligos diluted in 1X Buffer D were incubated on the cells for 30 min at 37°C. Finally, cells were washed with HSB (1x, 5 min), TBS (1x, 5 min), and 0.2X TBST (2x, 3 min), then mounted to slides using ProLong™ Gold Antifade Mountant with DNA Stain DAPI (Invitrogen, #P36935). Images were acquired on the FLUOVIEW FV3000 confocal microscope (Olympus). Analysis of the data was done with CellProfiler as described in the manufacturer's instructions.

Cell proliferation assay

$2.5x10^3$ cells were seeded into 96-well plates in 100µL of complete growth media. At 24, 48, 72, and 96 h post-seeding, 100µL CellTiter-Glo 2.0 (Promega, #G9242) was added to each well. Luminescence was measured with the BioTek Synergy H1 Plate Reader.

Scratch wound assay

Cells were seeded in 12-well plates to reach 90-100% confluency the following day, then incubated overnight at 37°C with 5% $CO_2$. The next morning, a scratch wound was made using a multichannel pipette equipped with P200 pipette tips. The cells were then washed 2x with PBS, and complete media was added to the wells. The rate of wound closure in each well was monitored via live cell imaging. While the cells were incubated at 37°C with 5% $CO_2$, brightfield images were taken every 30 min for 24 h on the Nikon Eclipse Ti2 microscope.

Cell cycle analysis

Wild-type PC-3 cells and SLK-KO PC-3 cells were seeded in 6-well plates to reach 80-100% confluency the following day, then incubated overnight at 37°C with 5% $CO_2$. The next morning, $1x10^6$ cells per replicate were harvested using trypsin, washed once with PBS, and pelleted in 15mL conical vials. Cells were fixed by slowly adding ice-cold 70% ethanol and incubating at 4°C for 1 h. The cells were washed 2x with PBS and resuspended in 0.5mL of FxCycle™ PI/RNase Staining Solution (Thermo Scientific, #F10797). Samples were incubated for 15-30 min at room temperature in the dark. The samples were then analyzed on the BD LSRII flow cytometer. The results were analyzed and plotted using FlowJo and GraphPad Prism 10.

*In vivo* tumor growth

All animal experimental procedures were approved by and performed in accordance with the guidelines of the Institutional Animal Care and Use Committee (protocol #18-031-03). Eight-week-old male NOD.Cg-Prkdc scid mice (Jackson Laboratory) were subcutaneously injected bilaterally with $1.2 \times 10^6$ PC-3 WT or PC-3 SLK-KO cells in DMEM (Corning)/Matrigel (Corning) (50/50). Tumor diameter was measured twice a week with a caliper. The experiment was terminated when the first animal reached maximum bilateral tumor size (1.5cm).

Cloning

c-Myc-HA-Cfa$^N$-FLAG was cloned into a pcDNA3.1 vector backbone using Gibson Assembly (New England Biolabs) following the manufacturer's instructions. The encoded construct was under the control of a CMV promoter. SLK-Short-3xFLAG-V5 was cloned by PCR amplifying SLK-Long-3xFLAG-V5 (VectorBuilder, VB250508-1503jms) using a forward primer annealing to the start of exon 14 and a reverse primer annealing to the end of exon 12 (see Table 1). Ligation of the amplicon yielded the SLK-Short-3xFLAG-V5 plasmid, lacking SLK exon 13. Plasmid sequences were verified by full plasmid sequencing (Plasmidsaurus). DNA sequences and corresponding amino acid sequences for the encoded proteins are provided below for each construct.

c-Myc-HA-CFA$^N$-FLAG

```
ATGCCCCTCAACGTTAGCTTCACCAACAGGAACTATGACCTCGACTACGACTCGGTGCAGCCGTATTTCTACTGCGACGAGGAGGAGAA
CTTCTACCAGCAGCAGCAGCAGAGCGAGCTGCAGCCCCCGGCGCCCAGCGAGGATATCTGGAAGAAATTCGAGCTGCTGCCCACCCCGC
CCCTGTCCCCTAGCCGCCGCTCCGGGCTCTGCTCGCCCTCCTACGTTGCGGTCACACCCTTCTCCCTTCGGGGAGACAACGACGGCGGT
GGCGGGAGCTTCTCCACGGCCGACCAGCTGGAGATGGTGACCGAGCTGCTGGGAGGAGACATGGTGAACCAGAGTTTCATCTGCGACCC
GGACGACGAGACCTTCATCAAAAACATCATCATCCAGGACTGTATGTGGAGCGGCTTCTCGGCCGCCGCCAAGCTCGTCTCAGAGAAGC
TGGCCTCCTACCAGGCTGCGCGCAAAGACAGCGGCAGCCCGAACCCCGCCCGCGGCCACAGCGTCTGCTCCACCTCCAGCTTGTACCTG
CAGGATCTGAGCGCCGCCGCCTCAGAGTGCATCGACCCCTCGGTGGTCTTCCCCTACCCTCTCAACGACAGCAGCTCGCCCAAGTCCTG
CGCCTCGCAAGACTCCAGCGCCTTCTCTCCGTCCTCGGATTCTCTGCTCTCCTCGACGGAGTCCTCCCCGCAGGGCAGCCCCGAGCCCC
TGGTGCTCCATGAGGAGACACCGCCCACCACCAGCAGCGACTCTGAGGAGGAACAAGAAGATGAGGAAGAAATCGATGTTGTTTCTGTG
GAAAAGAGGCAGGCTCCTGGCAAAAGGTCAGAGTCTGGATCACCTTCTGCTGGAGGCCACAGCAAACCTCCTCACAGCCCACTGGTCCT
CAAGAGGTGCCACGTCTCCACACATCAGCACAACTACGCAGCGCCTCCCTCCACTCGGAAGGACTATCCTGCTGCCAAGAGGGTCAAGT
TGGACAGTGTCAGAGTCCTGAGACAGATCAGCAACAACCGAAAATGCACCAGCCCCAGGTCCTCGGACACCGAGGAGAATGTCAAGAGG
CGAACACACAACGTCTTGGAGCGCCAGAGGAGGAACGAGCTAAAACGGAGCTTTTTTGCCCTGCGTGACCAGATCCCGGAGTTGGAAAA
CAATGAAAAGGCCCCCAAGGTAGTTATCCTTAAAAAAGCCACAGCATACATCCTGTCCGTCCAAGCAGAGGAGCAAAAGCTCATTTCTG
AAGAGGACTTGTTGCGGAAACGACGAGAACAGTTGAAACACAAACTTGAACAGCTACGGAACTCTTGTGCGGGTGGTTATCCGTATGAT
GTGCCGGATTATGCGTGCCTGTCTTACGACACAGAGATTCTGACCGTTGAATATGGATTCCTTCCTATCGGTAAGATCGTGGAGGAACG
GATTGAATGCACAGTCTATACGGTAGATAAAAATGGCTTTGTGTATACACAACCTATTGCTCAGTGGCATAACCGGGGAGAACAGGAAG
TTTTCGAATACTGCTTAGAAGACGGTTCGATTATCCGTGCAACGAAAGATCACAAATTTATGACGACCGACGGTCAGATGTTACCGATT
GATGAGATTTTCGAACGGGGGTTAGACCTGAAACAAGTTGATGGTTTGCCGAAAGGCGATTACAAGGATGACGACGATAAGTAA
```

```
MPLNVSFTNRNYDLDYDSVQPYFYCDEEENFYQQQQQSELQPPAPSEDIWKKFELLPTPPLSPSRRSGLCSPSYVAVTPFSLRGDNDGG
GGSFSTADQLEMVTELLGGDMVNQSFICDPDDETFIKNIIIQDCMWSGFSAAAKLVSEKLASYQAARKDSGSPNPARGHSVCSTSSLYL
QDLSAAASECIDPSVVFPYPLNDSSSPKSCASQDSSAFSPSSDSLLSSTESSPQGSPEPLVLHEETPPTTSSDSEEEQEDEEEIDVVSV
EKRQAPGKRSESGSPSAGGHSKPPHSPLVLKRCHVSTHQHNYAAPPSTRKDYPAAKRVKLDSVRVLRQISNNRKCTSPRSSDTEENVKR
RTHNVLERQRRNELKRSFFALRDQIPELENNEKAPKVVILKKATAYILSVQAEEQKLISEEDLLRKRREQLKHKLEQLRNSCAGGYPYD
VPDYACLSYDTEILTVEYGFLPIGKIVEERIECTVYTVDKNGFVYTQPIAQWHNRGEQEVFEYCLEDGSIIRATKDHKFMTTDGQMLPI
DEIFERGLDLKQVDGLPKGDYKDDDDK*
```

SLK-Short-3xFLAG-V5

ATGTCCTTCTTCAATTTCCGTAAGATCTTCAAGTTGGGGAGCGAGAAGAAGAAGAAGCAGTACGAACACGTGAAGAGGGACCTGAACCC
CGAAGACTTTTGGGAGATTATAGGAGAACTGGGCGACGGAGCCTTTGGGAAAGTGTACAAGGCCCAGAATAAAGAGACCAGTGTTTTAG
CTGCTGCAAAAGTGATTGACACTAAATCTGAAGAAGAACTTGAAGATTACATGGTAGAGATTGACATATTAGCATCTTGTGATCACCCA
AATATAGTCAAGCTTCTAGATGCCTTCTATTATGAGAACAATCTTTGGATCCTCATTGAATTTTGTGCAGGTGGAGCAGTAGATGCTGT
GATGCTTGAACTTGAGAGACCATTAACTGAGTCCCAAATACAAGTAGTTTGCAAGCAGACTTTAGATGCATTGAACTACTTACATGATA
ATAAGATCATCCACAGAGATCTGAAGGCTGGCAACATTCTCTTTACCTTAGATGGAGATATCAAATTGGCGGATTTTGGAGTATCAGCT
AAAAACACGAGGACAATTCAAAGAAGAGATTCCTTTATTGGTACACCATATTGGATGGCTCCTGAAGTAGTCATGTGTGAAACATCTAA
GGACAGACCCTATGACTACAAAGCTGATGTTTGGTCCCTGGGTATCACTTTAATAGAAATGGCTGAGATAGAACCACCTCATCATGAAT
TAAATCCAATGCGAGTGCTGCTAAAAATAGCAAAATCTGAGCCACCTACATTAGCACAGCCATCCAGATGGTCTTCAAATTTTAAGGAC
TTTCTAAAGAAATGCTTAGAAAAGAATGTGGATGCCAGGTGGACTACATCTCAGCTGCTGCAGCATCCCTTTGTTACTGTTGATTCCAA
CAAACCCATCCGAGAATTGATTGCAGAGGCGAAGGCTGAAGTAACAGAAGAAGTTGAAGATGGCAAAGAGGAAGATGAAGAGGAGGAAA
CAGAAAATTCTCTGCCAATACCTGCAAGTAAGCGTGCATCTTCTGACCTTAGTATCGCCAGCTCTGAAGAAGATAAACTTTCACAAAAT
GCTTGTATTTTGGAGTCTGTCTCAGAAAAAACAGAACGTAGTAACTCTGAAGATAAACTCAACAGCAAAATTCTTAATGAAAAACCCAC
CACTGATGAACCTGAAAAGGCTGTGGAGGATATTAATGAACATATTACCGATGCTCAGTTAGAAGCAATGACTGAACTCCATGACAGAA
CAGCAGTAATCAAGGAGAATGAAAGAGAGAAGAGGCCCAAGCTTGAAAATCTGCCTGACACAGAAGACCAAGAAACTGTGGACATTAAT
TCAGTCAGTGAAGGAAAAGAGAATAATATAATGATAACCTTAGAAACAAATATTGAACATAATCTAAAATCTGAGGAAGAAAAGGATCA
GGAAAAGCAACAGATGTTTGAAAATAAGCTTATAAAATCTGAAGAAATTAAAGATACTATTTTGCAAACAGTAGATTTAGTTTCTCAAG
AGACTGGAGAAAAAGAGGCAAATATTCAGGCAGTTGATAGTGAAGTTGGGCTTACAAAGGAAGACACCCAAGAGAAATTGGGGGAAGAC
GACAAAACTCAAAAAGATGTGATCAGCAATACAAGTGATGTGATAGGAACATGTGAGGCAGCAGATGTGGCTCAGAAAGTGGATGAAGA
CAGTGCTGAGGATACGCAGAGTAATGATGGGAAAGAAGTGGTCGAAGTAGGCCAGAAATTAATTAATAAGCCCATGGTGGGTCCTGAGG
CTGGTGGTACTAAGGAAGTTCCTATTAAAGAAATAGTTGAAATGAATGAAATAGAAGAAGGTAAAAATAAGGAACAAGCAATAAACAGT
TCAGAGAACATAATGGACATCAATGAGGAACCAGGAACAACTGAAGGTGAAGAAATCACTGAGTCAAGTAGCACTGAAGAAATGGAGGT
CAGAAGTGTGGTGGCTGATACTGACCAAAAGGCTTTAGGAAGTGAAGTTCAGGATGCTTCTAAAGTCACTACTCAGATAGATAAAGAGA
AAAAAGAAATTCCAGTGTCAATTAAAAAAGAGCCTGAAGTTACTGTAGTTTCACAGCCCACTGAACCTCAGCCTGTTCTAATACCCAGT
ATTAATATCAACTCTGACAGTGGAGAAAATAAAGAAGAAATAGGTTCTTTATCAAAAACTGAAACTATTCTGCCACCAGAATCTGAGAA
TCCAAAGGAAAATGATAATGATTCAGGCACTGGTTCCACTGCTGATACTAGCAGTATTGACTTGAATTTATCCATCTCTAGCTTTCTAA
GTAAAACTAAAGACAGTGGATCGATATCTTTACAAGAAACAAGAAGACAAAAGAAAACATTGAAGAAAACACGCAAATTTATTGTTGAT
GGTGTAGAAGTGAGTGTAACAACATCAAAGATAGTTACAGATAGTGATTCCAAAACTGAAGAATTGCGGTTTCTTAGACGTCAGGAACT
TCGGGAATTAAGATTTCTTCAGAAAGAAGAGCAAAGAGCCCAACAACAGCTCAATAGCAAACTACAGCAACAACGAGAACAAATTTTCC
GGCGCTTTGAGCAGGAAATGATGAGTAAAAAGCGACAATATGACCAGGAAATTGAGAATCTAGAAAAACAGCAGAAACAGACTATCGAA
CGCCTGGAACAAGAGCACACAAATCGCTTGCGAGATGAAGCCAAACGCATCAAAGGAGAACAAGAGAAAGAGTTGTCCAAATTTCAGAA
TATGCTGAAGAACCGAAAGAAGGAGGAACAAGAGTTTGTTCAGAAACAACAGCAAGAATTAGATGGCTCTCTGAAAAAGATCATCCAGC
AGCAGAAGGCAGAGTTAGCTAATATTGAGAGAGAGTGCCTGAATAACAAGCAACAGCTCATGAGAGCTCGAGAAGCTGCAATTTGGGAG
CTCGAAGAACGACACTTACAAGAAAAACACCAGCTGCTCAAACAGCAGCTTAAAGATCAGTATTTCATGCAAAGACATCAGCTACTTAA
GCGCCACGAGAAGGAAACAGAGCAAATGCAGCGTTACAATCAAAGACTTATTGAGGAATTGAAAAACAGACAGACTCAAGAAAGAGCAA
GACTGCCCAAGATTCAGCGCAGTGAAGCCAAGACTCGAATGGCCATGTTTAAGAAGAGTTTGAGAATTAACTCAACAGCCACACCAGAT
CAGGACCGTGATAAAATTAAACAGTTTGCTGCACAAGAAGAAAAGAGGCAGAAAAATGAGAGAATGGCTCAGCATCAGAAACATGAGAA
TCAAATGCGAGATCTTCAGTTGCAGTGTGAAGCCAATGTCCGCGAACTGCATCAGCTGCAGAATGAAAAATGCCACTTGTTGGTTGAGC
ATGAGACTCAGAAACTGAAGGAGTTAGATGAGGAACATAGCCAAGAATTAAAGGAGTGGAGAGAGAAATTGAGACCTAGGAAAAAGACA
CTGGAAGAAGAGTTTGCCAGGAAACTACAGGAACAGGAAGTATTCTTTAAAATGACTGGGGAGTCTGAATGCCTTAACCCATCAACACA
GAGCCGGATTTCCAAATTTTATCCTATTCCCAGCTTGCATTCCACCGGATCAGGCAGCGGCAGCGGCAGCGACTACAAAGACCATGACG
GTGATTATAAAGATCATGATATCGATTACAAGGATGACGATGACAAGGGCAGCGGCAGCGGCAGCGGTAAGCCTATCCCTAACCCTCTC
CTCGGTCTCGATTCTACGTAA

MSFFNFRKIFKLGSEKKKKQYEHVKRDLNPEDFWEIIGELGDGAFGKVYKAQNKETSVLAAAKVIDTKSEEELEDYMVEIDILASCDHP
NIVKLLDAFYYENNLWILIEFCAGGAVDAVMLELERPLTESQIQVVCKQTLDALNYLHDNKIIHRDLKAGNILFTLDGDIKLADFGVSA
KNTRTIQRRDSFIGTPYWMAPEVVMCETSKDRPYDYKADVWSLGITLIEMAEIEPPHHELNPMRVLLKIAKSEPPTLAQPSRWSSNFKD
FLKKCLEKNVDARWTTSQLLQHPFVTVDSNKPIRELIAEAKAEVTEEVEDGKEEDEEEETENSLPIPASKRASSDLSIASSEEDKLSQN
ACILESVSEKTERSNSEDKLNSKILNEKPTTDEPEKAVEDINEHITDAQLEAMTELHDRTAVIKENEREKRPKLENLPDTEDQETVDIN
SVSEGKENNIMITLETNIEHNLKSEEEKDQEKQQMFENKLIKSEEIKDTILQTVDLVSQETGEKEANIQAVDSEVGLTKEDTQEKLGED
DKTQKDVISNTSDVIGTCEAADVAQKVDEDSAEDTQSNDGKEVVEVGQKLINKPMVGPEAGGTKEVPIKEIVEMNEIEEGKNKEQAINS
SENIMDINEEPGTTEGEEITESSSTEEMEVRSVVADTDQKALGSEVQDASKVTTQIDKEKKEIPVSIKKEPEVTVVSQPTEPQPVLIPS
ININSDSGENKEEIGSLSKTETILPPESENPKENDNDSGTGSTADTSSIDLNLSISSFLSKTKDSGSISLQETRRQKKTLKKTRKFIVD
GVEVSVTTSKIVTDSDSKTEELRFLRRQELRELRFLQKEEQRAQQQLNSKLQQQREQIFRRFEQEMMSKKRQYDQEIENLEKQQKQTIE
RLEQEHTNRLRDEAKRIKGEQEKELSKFQNMLKNRKKEEQEFVQKQQQELDGSLKKIIQQQKAELANIERECLNNKQQLMRAREAAIWE
LEERHLQEKHQLLKQQLKDQYFMQRHQLLKRHEKETEQMQRYNQRLIEELKNRQTQERARLPKIQRSEAKTRMAMFKKSLRINSTATPD
QDRDKIKQFAAQEEKRQKNERMAQHQKHENQMRDLQLQCEANVRELHQLQNEKCHLLVEHETQKLKELDEEHSQELKEWREKLRPRKKT
LEEEFARKLQEQEVFFKMTGESECLNPSTQSRISKFYPIPSLHSTGSGSGSGSDYKDHDGDYKDHDIDYKDDDDKGSGSGSGKPIPNPL
LGLDST*

Commercial Plasmids

**Addgene:**
The following plasmids were purchased from Addgene and were made available by Dr. Linda Penn, University Health Network, Toronto, Canada.[11] The encoded constructs were under the control of a CMV promoter. Plasmid sequences were verified by full plasmid sequencing (Plasmidsaurus). DNA sequences and corresponding amino acid sequences for the encoded proteins are provided below for each construct.

c-Myc-V5-6xHis (Addgene, #176049)
ATGCCCCTCAACGTTAGCTTCACCAACAGGAACTATGACCTCGACTACGACTCGGTGCAGCCGTATTTCTACTGCGACGAGGAGGAGAA
CTTCTACCAGCAGCAGCAGCAGAGCGAGCTGCAGCCCCCGGCGCCCAGCGAGGATATCTGGAAGAAATTCGAGCTGCTGCCCACCCCGC
CCCTGTCCCCTAGCCGCCGCTCCGGGCTCTGCTCGCCCTCCTACGTTGCGGTCACACCATTCTCCCTTCGGGGAGACAACGACGGCGGT
GGCGGGAGCTTCTCCACGGCCGACCAGCTGGAGATGGTGACCGAGCTGCTGGGAGGAGACATGGTGAACCAGAGTTTCATCTGCGACCC
GGACGACGAGACCTTCATCAAAAACATCATCATCCAGGACTGTATGTGGAGCGGCTTCTCGGCCGCCGCCAAGCTCGTCTCAGAGAAGC
TGGCCTCCTACCAGGCTGCGCGCAAAGACAGCGGCAGCCCGAACCCCGCCCGCGGCCACAGCGTCTGCTCCACCTCCAGCTTGTACCTG
CAGGATCTGAGCGCCGCCGCCTCAGAGTGCATCGACCCCTCGGTGGTCTTCCCCTACCCTCTCAACGACAGCAGCTCGCCCAAGTCCTG
CGCCTCGCAAGACTCCAGCGCCTTCTCTCCGTCCTCGGATTCTCTGCTCTCCTCGACGGAGTCCTCCCCGCAGGGCAGCCCCGAGCCCC
TGGTGCTCCATGAGGAGACACCGCCCACCACCAGCAGCGACTCTGAGGAGGAACAAGAAGATGAGGAAGAAATCGATGTTGTTTCTGTG
GAAAAGAGGCAGGCTCCTGGCAAAAGGTCAGAGTCTGGATCACCTTCTGCTGGAGGCCACAGCAAACCTCCTCACAGCCCACTGGTCCT
CAAGAGGTGCCACGTCTCCACACATCAGCACAACTACGCAGCGCCTCCCTCCACTCGGAAGGACTATCCTGCTGCCAAGAGGGTCAAGT
TGGACAGTGTCAGAGTCCTGAGACAGATCAGCAACAACCGAAAATGCACCAGCCCCAGGTCCTCGGACACCGAGGAGAATGTCAAGAGG
CGAACACACAACGTCTTGGAGCGCCAGAGGAGGAACGAGCTAAAACGGAGCTTTTTTGCCCTGCGTGACCAGATCCCGGAGTTGGAAAA
CAATGAAAAGGCCCCCAAGGTAGTTATCCTTAAAAAAGCCACAGCATACATCCTGTCCGTCCAAGCAGAGGAGCAAAAGCTCATTTCTG
AAGAGGACTTGTTGCGGAAACGACGAGAACAGTTGAAACACAAACTTGAACAGCTACGGAACTCTTGTGCGGGTCTAGAGGGCCCGCGG
TTCGAAGGTAAGCCTATCCCTAACCCTCTCCTCGGTCTCGATTCTACGCGTACCGGTCATCATCACCATCACCATTGA

MPLNVSFTNRNYDLDYDSVQPYFYCDEEENFYQQQQQSELQPPAPSEDIWKKFELLPTPPLSPSRRSGLCSPSYVAVTPFSLRGDNDGG
GGSFSTADQLEMVTELLGGDMVNQSFICDPDDETFIKNIIIQDCMWSGFSAAAKLVSEKLASYQAARKDSGSPNPARGHSVCSTSSLYL
QDLSAAASECIDPSVVFPYPLNDSSSPKSCASQDSSAFSPSSDSLLSSTESSPQGSPEPLVLHEETPPTTSSDSEEEQEDEEEIDVVSV
EKRQAPGKRSESGSPSAGGHSKPPHSPLVLKRCHVSTHQHNYAAPPSTRKDYPAAKRVKLDSVRVLRQISNNRKCTSPRSSDTEENVKR
RTHNVLERQRRNELKRSFFALRDQIPELENNEKAPKVVILKKATAYILSVQAEEQKLISEEDLLRKRREQLKHKLEQLRNSCAGLEGPR
FEGKPIPNPLLGLDSTRTGHHHHHH*

c-Myc-V5-6xHis [ΔMyc-box 0] (Addgene, #176050)
ATGCCCCTCAACGTTAGCTTCACCAACAGGAACTATGACCTCGACCAGCAGCAGCAGAGCGAGCTGCAGCCCCCGGCGCCCAGCGAGGA
TATCTGGAAGAAATTCGAGCTGCTGCCCACCCCGCCCCTGTCCCCTAGCCGCCGCTCCGGGCTCTGCTCGCCCTCCTACGTTGCGGTCA
CACCCTTCTCCCTTCGGGGAGACAACGACGGCGGTGGCGGGAGCTTCTCCACGGCCGACCAGCTGGAGATGGTGACCGAGCTGCTGGGA
GGAGACATGGTGAACCAGAGTTTCATCTGCGACCCGGACGACGAGACCTTCATCAAAAACATCATCATCCAGGACTGTATGTGGAGCGG
CTTCTCGGCCGCCGCCAAGCTCGTCTCAGAGAAGCTGGCCTCCTACCAGGCTGCGCGCAAAGACAGCGGCAGCCCGAACCCCGCCCGCG
GCCACAGCGTCTGCTCCACCTCCAGCTTGTACCTGCAGGATCTGAGCGCCGCCGCCTCAGAGTGCATCGACCCCTCGGTGGTCTTCCCC
TACCCTCTCAACGACAGCAGCTCGCCCAAGTCCTGCGCCTCGCAAGACTCCAGCGCCTTCTCTCCGTCCTCGGATTCTCTGCTCTCCTC
GACGGAGTCCTCCCCGCAGGGCAGCCCCGAGCCCCTGGTGCTCCATGAGGAGACACCGCCCACCACCAGCAGCGACTCTGAGGAGGAAC
AAGAAGATGAGGAAGAAATCGATGTTGTTTCTGTGGAAAAGAGGCAGGCTCCTGGCAAAAGGTCAGAGTCTGGATCACCTTCTGCTGGA
GGCCACAGCAAACCTCCTCACAGCCCACTGGTCCTCAAGAGGTGCCACGTCTCCACACATCAGCACAACTACGCAGCGCCTCCCTCCAC
TCGGAAGGACTATCCTGCTGCCAAGAGGGTCAAGTTGGACAGTGTCAGAGTCCTGAGACAGATCAGCAACAACCGAAAATGCACCAGCC
CCAGGTCCTCGGACACCGAGGAGAATGTCAAGAGGCGAACACACAACGTCTTGGAGCGCCAGAGGAGGAACGAGCTAAAACGGAGCTTT
TTTGCCCTGCGTGACCAGATCCCGGAGTTGGAAAACAATGAAAAGGCCCCCAAGGTAGTTATCCTTAAAAAAGCCACAGCATACATCCT
GTCCGTCCAAGCAGAGGAGCAAAAGCTCATTTCTGAAGAGGACTTGTTGCGGAAACGACGAGAACAGTTGAAACACAAACTTGAACAGC
TACGGAACTCTTGTGCGGGTCTAGAGGGCCCGCGGTTCGAAGGTAAGCCTATCCCTAACCCTCTCCTCGGTCTCGATTCTACGCGTACC
GGTCATCATCACCATCACCATTGA

MPLNVSFTNRNYDLDQQQQSELQPPAPSEDIWKKFELLPTPPLSPSRRSGLCSPSYVAVTPFSLRGDNDGGGGSFSTADQLEMVTELLG
GDMVNQSFICDPDDETFIKNIIIQDCMWSGFSAAAKLVSEKLASYQAARKDSGSPNPARGHSVCSTSSLYLQDLSAAASECIDPSVVFP
YPLNDSSSPKSCASQDSSAFSPSSDSLLSSTESSPQGSPEPLVLHEETPPTTSSDSEEEQEDEEEIDVVSVEKRQAPGKRSESGSPSAG
GHSKPPHSPLVLKRCHVSTHQHNYAAPPSTRKDYPAAKRVKLDSVRVLRQISNNRKCTSPRSSDTEENVKRRTHNVLERQRRNELKRSF
FALRDQIPELENNEKAPKVVILKKATAYILSVQAEEQKLISEEDLLRKRREQLKHKLEQLRNSCAGLEGPRFEGKPIPNPLLGLDSTRT
GHHHHHH*

c-Myc-V5-6xHis [ΔMyc-box I] (Addgene, #176051)

ATGCCCCTCAACGTTAGCTTCACCAACAGGAACTATGACCTCGACTACGACTCGGTGCAGCCGTATTTCTACTGCGACGAGGAGGAGAA
CTTCTACCAGCAGCAGCAGCAGAGCGAGCTGCAGCCCCCGGCGAGCCGCCGCTCCGGGCTCTGCTCGCCCTCCTACGTTGCGGTCACAC
CCTTCTCCCTTCGGGGAGACAACGACGGCGGTGGCGGGAGCTTCTCCACGGCCGACCAGCTGGAGATGGTGACCGAGCTGCTGGGAGGA
GACATGGTGAACCAGAGTTTCATCTGCGACCCGGACGACGAGACCTTCATCAAAAACATCATCATCCAGGACTGTATGTGGAGCGGCTT
CTCGGCCGCCGCCAAGCTCGTCTCAGAGAAGCTGGCCTCCTACCAGGCTGCGCGCAAAGACAGCGGCAGCCCGAACCCCGCCCGCGGCC
ACAGCGTCTGCTCCACCTCCAGCTTGTACCTGCAGGATCTGAGCGCCGCCGCCTCAGAGTGCATCGACCCCTCGGTGGTCTTCCCCTAC
CCTCTCAACGACAGCAGCTCGCCCAAGTCCTGCGCCTCGCAAGACTCCAGCGCCTTCTCTCCGTCCTCGGATTCTCTGCTCTCCTCGAC
GGAGTCCTCCCCGCAGGGCAGCCCCGAGCCCCTGGTGCTCCATGAGGAGACACCGCCCACCACCAGCAGCGACTCTGAGGAGGAACAAG
AAGATGAGGAAGAAATCGATGTTGTTTCTGTGGAAAAGAGGCAGGCTCCTGGCAAAAGGTCAGAGTCTGGATCACCTTCTGCTGGAGGC
CACAGCAAACCTCCTCACAGCCCACTGGTCCTCAAGAGGTGCCACGTCTCCACACATCAGCACAACTACGCAGCGCCTCCCTCCACTCG
GAAGGACTATCCTGCTGCCAAGAGGGTCAAGTTGGACAGTGTCAGAGTCCTGAGACAGATCAGCAACAACCGAAAATGCACCAGCCCCA
GGTCCTCGGACACCGAGGAGAATGTCAAGAGGCGAACACACAACGTCTTGGAGCGCCAGAGGAGGAACGAGCTAAAACGGAGCTTTTTT
GCCCTGCGTGACCAGATCCCGGAGTTGGAAAACAATGAAAAGGCCCCCAAGGTAGTTATCCTTAAAAAAGCCACAGCATACATCCTGTC
CGTCCAAGCAGAGGAGCAAAAGCTCATTTCTGAAGAGGACTTGTTGCGGAAACGACGAGAACAGTTGAAACACAAACTTGAACAGCTAC
GGAACTCTTGTGCGGGTCTAGAGGGCCCGCGGTTCGAAGGTAAGCCTATCCCTAACCCTCTCCTCGGTCTCGATTCTACGCGTACCGGT
CATCATCACCATCACCATTGA

MPLNVSFTNRNYDLDYDSVQPYFYCDEEENFYQQQQQSELQPPASRRSGLCSPSYVAVTPFSLRGDNDGGGGSFSTADQLEMVTELLGG
DMVNQSFICDPDDETFIKNIIIQDCMWSGFSAAAKLVSEKLASYQAARKDSGSPNPARGHSVCSTSSLYLQDLSAAASECIDPSVVFPY
PLNDSSSPKSCASQDSSAFSPSSDSLLSSTESSPQGSPEPLVLHEETPPTTSSDSEEEQEDEEEIDVVSVEKRQAPGKRSESGSPSAGG
HSKPPHSPLVLKRCHVSTHQHNYAAPPSTRKDYPAAKRVKLDSVRVLRQISNNRKCTSPRSSDTEENVKRRTHNVLERQRRNELKRSFF
ALRDQIPELENNEKAPKVVILKKATAYILSVQAEEQKLISEEDLLRKRREQLKHKLEQLRNSCAGLEGPRFEGKPIPNPLLGLDSTRTG
HHHHHH*

c-Myc-V5-6xHis [ΔMyc-box II] (Addgene, #176052)

ATGCCCCTCAACGTTAGCTTCACCAACAGGAACTATGACCTCGACTACGACTCGGTGCAGCCGTATTTCTACTGCGACGAGGAGGAGAA
CTTCTACCAGCAGCAGCAGCAGAGCGAGCTGCAGCCCCCGGCGCCCAGCGAGGATATCTGGAAGAAATTCGAGCTGCTGCCCACCCCGC
CCCTGTCCCCTAGCCGCCGCTCCGGGCTCTGCTCGCCCTCCTACGTTGCGGTCACACCCTTCTCCCTTCGGGGAGACAACGACGGCGGT
GGCGGGAGCTTCTCCACGGCCGACCAGCTGGAGATGGTGACCGAGCTGCTGGGAGGAGACATGGTGAACCAGAGTTTCATCTGCGACCC
GGACGACGAGACCTTCATCAAAAACCTCGTCTCAGAGAAGCTGGCCTCCTACCAGGCTGCGCGCAAAGACAGCGGCAGCCCGAACCCCG
CCCGCGGCCACAGCGTCTGCTCCACCTCCAGCTTGTACCTGCAGGATCTGAGCGCCGCCGCCTCAGAGTGCATCGACCCCTCGGTGGTC
TTCCCCTACCCTCTCAACGACAGCAGCTCGCCCAAGTCCTGCGCCTCGCAAGACTCCAGCGCCTTCTCTCCGTCCTCGGATTCTCTGCT
CTCCTCGACGGAGTCCTCCCCGCAGGGCAGCCCCGAGCCCCTGGTGCTCCATGAGGAGACACCGCCCACCACCAGCAGCGACTCTGAGG
AGGAACAAGAAGATGAGGAAGAAATCGATGTTGTTTCTGTGGAAAAGAGGCAGGCTCCTGGCAAAAGGTCAGAGTCTGGATCACCTTCT
GCTGGAGGCCACAGCAAACCTCCTCACAGCCCACTGGTCCTCAAGAGGTGCCACGTCTCCACACATCAGCACAACTACGCAGCGCCTCC
CTCCACTCGGAAGGACTATCCTGCTGCCAAGAGGGTCAAGTTGGACAGTGTCAGAGTCCTGAGACAGATCAGCAACAACCGAAAATGCA
CCAGCCCCAGGTCCTCGGACACCGAGGAGAATGTCAAGAGGCGAACACACAACGTCTTGGAGCGCCAGAGGAGGAACGAGCTAAAACGG
AGCTTTTTTGCCCTGCGTGACCAGATCCCGGAGTTGGAAAACAATGAAAAGGCCCCCAAGGTAGTTATCCTTAAAAAAGCCACAGCATA
CATCCTGTCCGTCCAAGCAGAGGAGCAAAAGCTCATTTCTGAAGAGGACTTGTTGCGGAAACGACGAGAACAGTTGAAACACAAACTTG
AACAGCTACGGAACTCTTGTGCGGGTCTAGAGGGCCCGCGGTTCGAAGGTAAGCCTATCCCTAACCCTCTCCTCGGTCTCGATTCTACG
CGTACCGGTCATCATCACCATCACCATTGA
MPLNVSFTNRNYDLDYDSVQPYFYCDEEENFYQQQQQSELQPPAPSEDIWKKFELLPTPPLSPSRRSGLCSPSYVAVTPFSLRGDNDGG
GGSFSTADQLEMVTELLGGDMVNQSFICDPDDETFIKNLVSEKLASYQAARKDSGSPNPARGHSVCSTSSLYLQDLSAAASECIDPSVV
FPYPLNDSSSPKSCASQDSSAFSPSSDSLLSSTESSPQGSPEPLVLHEETPPTTSSDSEEEQEDEEEIDVVSVEKRQAPGKRSESGSPS
AGGHSKPPHSPLVLKRCHVSTHQHNYAAPPSTRKDYPAAKRVKLDSVRVLRQISNNRKCTSPRSSDTEENVKRRTHNVLERQRRNELKR
SFFALRDQIPELENNEKAPKVVILKKATAYILSVQAEEQKLISEEDLLRKRREQLKHKLEQLRNSCAGLEGPRFEGKPIPNPLLGLDST
RTGHHHHHH*

c-Myc-V5-6xHis [ΔMyc-box IIIa] (Addgene, #176053)

ATGCCCCTCAACGTTAGCTTCACCAACAGGAACTATGACCTCGACTACGACTCGGTGCAGCCGTATTTCTACTGCGACGAGGAGGAGAA
CTTCTACCAGCAGCAGCAGCAGAGCGAGCTGCAGCCCCCGGCGCCCAGCGAGGATATCTGGAAGAAATTCGAGCTGCTGCCCACCCCGC
CCCTGTCCCCTAGCCGCCGCTCCGGGCTCTGCTCGCCCTCCTACGTTGCGGTCACACCCTTCTCCCTTCGGGGAGACAACGACGGCGGT
GGCGGGAGCTTCTCCACGGCCGACCAGCTGGAGATGGTGACCGAGCTGCTGGGAGGAGACATGGTGAACCAGAGTTTCATCTGCGACCC
GGACGACGAGACCTTCATCAAAAACATCATCATCCAGGACTGTATGTGGAGCGGCTTCTCGGCCGCCGCCAAGCTCGTCTCAGAGAAGC
TGGCCTCCTACCAGGCTGCGCGCAAAGACAGCGGCAGCCCGAACCCCGCCCGCGGCCACAGCGTCTGCTCCACCTCCAGCTTGTACCTG
CAGGATCTGAGCGCCGCCGCCTCAGAGAACGACAGCAGCTCGCCCAAGTCCTGCGCCTCGCAAGACTCCAGCGCCTTCTCTCCGTCCTC
GGATTCTCTGCTCTCCTCGACGGAGTCCTCCCCGCAGGGCAGCCCCGAGCCCCTGGTGCTCCATGAGGAGACACCGCCCACCACCAGCA
GCGACTCTGAGGAGGAACAAGAAGATGAGGAAGAAATCGATGTTGTTTCTGTGGAAAAGAGGCAGGCTCCTGGCAAAAGGTCAGAGTCT
GGATCACCTTCTGCTGGAGGCCACAGCAAACCTCCTCACAGCCCACTGGTCCTCAAGAGGTGCCACGTCTCCACACATCAGCACAACTA
CGCAGCGCCTCCCTCCACTCGGAAGGACTATCCTGCTGCCAAGAGGGTCAAGTTGGACAGTGTCAGAGTCCTGAGACAGATCAGCAACA

ACCGAAAATGCACCAGCCCCAGGTCCTCGGACACCGAGGAGAATGTCAAGAGGCGAACACACAACGTCTTGGAGCGCCAGAGGAGGAAC
GAGCTAAAACGGAGCTTTTTTGCCCTGCGTGACCAGATCCCGGAGTTGGAAAACAATGAAAAGGCCCCCAAGGTAGTTATCCTTAAAAA
AGCCACAGCATACATCCTGTCCGTCCAAGCAGAGGAGCAAAAGCTCATTTCTGAAGAGGACTTGTTGCGGAAACGACGAGAACAGTTGA
AACACAAACTTGAACAGCTACGGAACTCTTGTGCGGGTCTAGAGGGCCCGCGGTTCGAAGGTAAGCCTATCCCTAACCCTCTCCTCGGT
CTCGATTCTACGCGTACCGGTCATCATCACCATCACCATTGA

MPLNVSFTNRNYDLDYDSVQPYFYCDEEENFYQQQQQSELQPPAPSEDIWKKFELLPTPPLSPSRRSGLCSPSYVAVTPFSLRGDNDGG
GGSFSTADQLEMVTELLGGDMVNQSFICDPDDETFIKNIIIQDCMWSGFSAAAKLVSEKLASYQAARKDSGSPNPARGHSVCSTSSLYL
QDLSAAASENDSSSPKSCASQDSSAFSPSSDSLLSSTESSPQGSPEPLVLHEETPPTTSSDSEEEQEDEEEIDVVSVEKRQAPGKRSES
GSPSAGGHSKPPHSPLVLKRCHVSTHQHNYAAPPSTRKDYPAAKRVKLDSVRVLRQISNNRKCTSPRSSDTEENVKRRTHNVLERQRRN
ELKRSFFALRDQIPELENNEKAPKVVILKKATAYILSVQAEEQKLISEEDLLRKRREQLKHKLEQLRNSCAGLEGPRFEGKPIPNPLLG
LDSTRTGHHHHHH*

#### c-Myc-V5-6xHis [ΔMyc-box IIIb] (Addgene, #176054)

ATGCCCCTCAACGTTAGCTTCACCAACAGGAACTATGACCTCGACTACGACTCGGTGCAGCCGTATTTCTACTGCGACGAGGAGGAGAA
CTTCTACCAGCAGCAGCAGCAGAGCGAGCTGCAGCCCCCGGCGCCCAGCGAGGATATCTGGAAGAAATTCGAGCTGCTGCCCACCCCGC
CCCTGTCCCCTAGCCGCCGCTCCGGGCTCTGCTCGCCCTCCTACGTTGCGGTCACACCCTTCTCCCTTCGGGGAGACAACGACGGCGGT
GGCGGGAGCTTCTCCACGGCCGACCAGCTGGAGATGGTGACCGAGCTGCTGGGAGGAGACATGGTGAACCAGAGTTTCATCTGCGACCC
GGACGACGAGACCTTCATCAAAAACATCATCATCCAGGACTGTATGTGGAGCGGCTTCTCGGCCGCCGCCAAGCTCGTCTCAGAGAAGC
TGGCCTCCTACCAGGCTGCGCGCAAAGACAGCGGCAGCCCGAACCCCGCCCGCGGCCACAGCGTCTGCTCCACCTCCAGCTTGTACCTG
CAGGATCTGAGCGCCGCCGCCTCAGAGTGCATCGACCCCTCGGTGGTCTTCCCCTACCCTCTCAACGACAGCAGCTCGCCCAAGTCCTG
CGCCTCGCAAGACTCCAGCGCCTTCTCTCCGTCCTCGGATTCTCTGCTCTCCTCGACGGAGTCCTCCCCGCAGGGCAGCCCCGAGCCCC
TGGTGCTCCATGAGGAGACACCGCCCACCACCAGCAGCGACTCTGAGGAGGAACAAGAAGATCAGGCTCCTGGCAAAAGGTCAGAGTCT
GGATCACCTTCTGCTGGAGGCCACAGCAAACCTCCTCACAGCCCACTGGTCCTCAAGAGGTGCCACGTCTCCACACATCAGCACAACTA
CGCAGCGCCTCCCTCCACTCGGAAGGACTATCCTGCTGCCAAGAGGGTCAAGTTGGACAGTGTCAGAGTCCTGAGACAGATCAGCAACA
ACCGAAAATGCACCAGCCCCAGGTCCTCGGACACCGAGGAGAATGTCAAGAGGCGAACACACAACGTCTTGGAGCGCCAGAGGAGGAAC
GAGCTAAAACGGAGCTTTTTTGCCCTGCGTGACCAGATCCCGGAGTTGGAAAACAATGAAAAGGCCCCCAAGGTAGTTATCCTTAAAAA
AGCCACAGCATACATCCTGTCCGTCCAAGCAGAGGAGCAAAAGCTCATTTCTGAAGAGGACTTGTTGCGGAAACGACGAGAACAGTTGA
AACACAAACTTGAACAGCTACGGAACTCTTGTGCGGGTCTAGAGGGCCCGCGGTTCGAAGGTAAGCCTATCCCTAACCCTCTCCTCGGT
CTCGATTCTACGCGTACCGGTCATCATCACCATCACCATTGA

MPLNVSFTNRNYDLDYDSVQPYFYCDEEENFYQQQQQSELQPPAPSEDIWKKFELLPTPPLSPSRRSGLCSPSYVAVTPFSLRGDNDGG
GGSFSTADQLEMVTELLGGDMVNQSFICDPDDETFIKNIIIQDCMWSGFSAAAKLVSEKLASYQAARKDSGSPNPARGHSVCSTSSLYL
QDLSAAASECIDPSVVFPYPLNDSSSPKSCASQDSSAFSPSSDSLLSSTESSPQGSPEPLVLHEETPPTTSSDSEEEQEDQAPGKRSES
GSPSAGGHSKPPHSPLVLKRCHVSTHQHNYAAPPSTRKDYPAAKRVKLDSVRVLRQISNNRKCTSPRSSDTEENVKRRTHNVLERQRRN
ELKRSFFALRDQIPELENNEKAPKVVILKKATAYILSVQAEEQKLISEEDLLRKRREQLKHKLEQLRNSCAGLEGPRFEGKPIPNPLLG
LDSTRTGHHHHHH*

#### c-Myc-V5-6xHis [ΔMyc-box IV] (#176055)

ATGCCCCTCAACGTTAGCTTCACCAACAGGAACTATGACCTCGACTACGACTCGGTGCAGCCGTATTTCTACTGCGACGAGGAGGAGAA
CTTCTACCAGCAGCAGCAGCAGAGCGAGCTGCAGCCCCCGGCGCCCAGCGAGGATATCTGGAAGAAATTCGAGCTGCTGCCCACCCCGC
CCCTGTCCCCTAGCCGCCGCTCCGGGCTCTGCTCGCCCTCCTACGTTGCGGTCACACCCTTCTCCCTTCGGGGAGACAACGACGGCGGT
GGCGGGAGCTTCTCCACGGCCGACCAGCTGGAGATGGTGACCGAGCTGCTGGGAGGAGACATGGTGAACCAGAGTTTCATCTGCGACCC
GGACGACGAGACCTTCATCAAAAACATCATCATCCAGGACTGTATGTGGAGCGGCTTCTCGGCCGCCGCCAAGCTCGTCTCAGAGAAGC
TGGCCTCCTACCAGGCTGCGCGCAAAGACAGCGGCAGCCCGAACCCCGCCCGCGGCCACAGCGTCTGCTCCACCTCCAGCTTGTACCTG
CAGGATCTGAGCGCCGCCGCCTCAGAGTGCATCGACCCCTCGGTGGTCTTCCCCTACCCTCTCAACGACAGCAGCTCGCCCAAGTCCTG
CGCCTCGCAAGACTCCAGCGCCTTCTCTCCGTCCTCGGATTCTCTGCTCTCCTCGACGGAGTCCTCCCCGCAGGGCAGCCCCGAGCCCC
TGGTGCTCCATGAGGAGACACCGCCCACCACCAGCAGCGACTCTGAGGAGGAACAAGAAGATGAGGAAGAAATCGATGTTGTTTCTGTG
GAAAAGAGGCAGGCTCCTGGCAAAAGGTCAGAGTCTGGATCACCTTCTGCTGGAGGCCACAGCAAACCTCCTCACAGCCCACTGGTCCT
CAAGAGGTGCCACGTCTCCGTCAAGTTGGACAGTGTCAGAGTCCTGAGACAGATCAGCAACAACCGAAAATGCACCAGCCCCAGGTCCT
CGGACACCGAGGAGAATGTCAAGAGGCGAACACACAACGTCTTGGAGCGCCAGAGGAGGAACGAGCTAAAACGGAGCTTTTTTGCCCTG
CGTGACCAGATCCCGGAGTTGGAAAACAATGAAAAGGCCCCCAAGGTAGTTATCCTTAAAAAAGCCACAGCATACATCCTGTCCGTCCA
AGCAGAGGAGCAAAAGCTCATTTCTGAAGAGGACTTGTTGCGGAAACGACGAGAACAGTTGAAACACAAACTTGAACAGCTACGGAACT
CTTGTGCGGGTCTAGAGGGCCCGCGGTTCGAAGGTAAGCCTATCCCTAACCCTCTCCTCGGTCTCGATTCTACGCGTACCGGTCATCAT
CACCATCACCATTGA

MPLNVSFTNRNYDLDYDSVQPYFYCDEEENFYQQQQQSELQPPAPSEDIWKKFELLPTPPLSPSRRSGLCSPSYVAVTPFSLRGDNDGG
GGSFSTADQLEMVTELLGGDMVNQSFICDPDDETFIKNIIIQDCMWSGFSAAAKLVSEKLASYQAARKDSGSPNPARGHSVCSTSSLYL
QDLSAAASECIDPSVVFPYPLNDSSSPKSCASQDSSAFSPSSDSLLSSTESSPQGSPEPLVLHEETPPTTSSDSEEEQEDEEEIDVVSV
EKRQAPGKRSESGSPSAGGHSKPPHSPLVLKRCHVSVKLDSVRVLRQISNNRKCTSPRSSDTEENVKRRTHNVLERQRRNELKRSFFAL

RDQIPELENNEKAPKVVILKKATAYILSVQAEEQKLISEEDLLRKRREQLKHKLEQLRNSCAGLEGPRFEGKPIPNPLLGLDSTRTGHH
HHHH*

pSpCas9(BB)-2A-GFP (PX458) (#48138)

gRNA sequences (see Table 1) were cloned into this plasmid backbone to generate CRISPR knockout cell lines.

**Vector Builder:**
The following plasmids were purchased from VectorBuilder. The encoded constructs were under the control of an EFS promoter. Plasmid sequences were verified by full plasmid sequencing (Plasmidsaurus). DNA sequences and corresponding amino acid sequences for the encoded proteins are provided below for each construct.

SLK-Long-3xFLAG-V5 (VB250508-1503jms)
ATGTCCTTCTTCAATTTCCGTAAGATCTTCAAGTTGGGGAGCGAGAAGAAGAAGAAGCAGTACGAACACGTGAAGAGGGACCTGAACCC
CGAAGACTTTTGGGAGATTATAGGAGAACTGGGCGACGGAGCCTTTGGGAAAGTGTACAAGGCCCAGAATAAAGAGACCAGTGTTTTAG
CTGCTGCAAAAGTGATTGACACTAAATCTGAAGAAGAACTTGAAGATTACATGGTAGAGATTGACATATTAGCATCTTGTGATCACCCA
AATATAGTCAAGCTTCTAGATGCCTTCTATTATGAGAACAATCTTTGGATCCTCATTGAATTTTGTGCAGGTGGAGCAGTAGATGCTGT
GATGCTTGAACTTGAGAGACCATTAACTGAGTCCCAAATACAAGTAGTTTGCAAGCAGACTTTAGATGCATTGAACTACTTACATGATA
ATAAGATCATCCACAGAGATCTGAAGGCTGGCAACATTCTCTTTACCTTAGATGGAGATATCAAATTGGCGGATTTTGGAGTATCAGCT
AAAAACACGAGGACAATTCAAAGAAGAGATTCCTTTATTGGTACACCATATTGGATGGCTCCTGAAGTAGTCATGTGTGAAACATCTAA
GGACAGACCCTATGACTACAAAGCTGATGTTTGGTCCCTGGGTATCACTTTAATAGAAATGGCTGAGATAGAACCACCTCATCATGAAT
TAAATCCAATGCGAGTGCTGCTAAAAATAGCAAAATCTGAGCCACCTACATTAGCACAGCCATCCAGATGGTCTTCAAATTTTAAGGAC
TTTCTAAAGAAATGCTTAGAAAAGAATGTGGATGCCAGGTGGACTACATCTCAGCTGCTGCAGCATCCCTTTGTTACTGTTGATTCCAA
CAAACCCATCCGAGAATTGATTGCAGAGGCGAAGGCTGAAGTAACAGAAGAAGTTGAAGATGGCAAAGAGGAAGATGAAGAGGAGGAAA
CAGAAAATTCTCTGCCAATACCTGCAAGTAAGCGTGCATCTTCTGACCTTAGTATCGCCAGCTCTGAAGAAGATAAACTTTCACAAAAT
GCTTGTATTTTGGAGTCTGTCTCAGAAAAAACAGAACGTAGTAACTCTGAAGATAAACTCAACAGCAAAATTCTTAATGAAAAACCCAC
CACTGATGAACCTGAAAAGGCTGTGGAGGATATTAATGAACATATTACCGATGCTCAGTTAGAAGCAATGACTGAACTCCATGACAGAA
CAGCAGTAATCAAGGAGAATGAAAGAGAGAAGAGGCCCAAGCTTGAAAATCTGCCTGACACAGAAGACCAAGAAACTGTGGACATTAAT
TCAGTCAGTGAAGGAAAAGAGAATAATATAATGATAACCTTAGAAACAAATATTGAACATAATCTAAAATCTGAGGAAGAAAAGGATCA
GGAAAAGCAACAGATGTTTGAAAATAAGCTTATAAAATCTGAAGAAATTAAAGATACTATTTTGCAAACAGTAGATTTAGTTTCTCAAG
AGACTGGAGAAAAAGAGGCAAATATTCAGGCAGTTGATAGTGAAGTTGGGCTTACAAAGGAAGACACCCAAGAGAAATTGGGGGAAGAC
GACAAAACTCAAAAAGATGTGATCAGCAATACAAGTGATGTGATAGGAACATGTGAGGCAGCAGATGTGGCTCAGAAAGTGGATGAAGA
CAGTGCTGAGGATACGCAGAGTAATGATGGGAAAGAAGTGGTCGAAGTAGGCCAGAAATTAATTAATAAGCCCATGGTGGGTCCTGAGG
CTGGTGGTACTAAGGAAGTTCCTATTAAAGAAATAGTTGAAATGAATGAAATAGAAGAAGGTAAAAATAAGGAACAAGCAATAAACAGT
TCAGAGAACATAATGGACATCAATGAGGAACCAGGAACAACTGAAGGTGAAGAAATCACTGAGTCAAGTAGCACTGAAGAAATGGAGGT
CAGAAGTGTGGTGGCTGATACTGACCAAAAGGCTTTAGGAAGTGAAGTTCAGGATGCTTCTAAAGTCACTACTCAGATAGATAAAGAGA
AAAAAGAAATTCCAGTGTCAATTAAAAAAGAGCCTGAAGTTACTGTAGTTTCACAGCCCACTGAACCTCAGCCTGTTCTAATACCCAGT
ATTAATATCAACTCTGACAGTGGAGAAAATAAAGAAGAAATAGGTTCTTTATCAAAAACTGAAACTATTCTGCCACCAGAATCTGAGAA
TCCAAAGGAAAATGATAATGATTCAGGCACTGGTTCCACTGCTGATACTAGCAGTATTGACTTGAATTTATCCATCTCTAGCTTTCTAA
GTAAAACTAAAGACAGTGGATCGATATCTTTACAAGAAACAAGAAGACAAAAGAAAACATTGAAGAAAACACGCAAATTTATTGTTGAT
GGTGTAGAAGTGAGTGTAACAACATCAAAGATAGTTACAGATAGTGATTCCAAAACTGAAGAATTGCGGTTTCTTAGACGTCAGGAACT
TCGGGAATTAAGATTTCTTCAGAAAGAAGAGCAAAGAGCCCAACAACAGCTCAATAGCAAACTACAGCAACAACGAGAACAAATTTTCC
GGCGCTTTGAGCAGGAAATGATGAGTAAAAAGCGACAATATGACCAGGAAATTGAGAATCTAGAAAAACAGCAGAAACAGACTATCGAA
CGCCTGGAACAAGAGCACACAAATCGCTTGCGAGATGAAGCCAAACGCATCAAAGGAGAACAAGAGAAAGAGTTGTCCAAATTTCAGAA
TATGCTGAAGAACCGAAAGAAGGAGGTTATAAATGAAGTGGAGAAAGCACCCAAAGAGCTGAGAAAAGAGCTCATGAAACGCAGGAAAG
AGGAGCTTGCACAAAGCCAGCATGCTCAGGAACAAGAGTTTGTTCAGAAACAACAGCAAGAATTAGATGGCTCTCTGAAAAAGATCATC
CAGCAGCAGAAGGCAGAGTTAGCTAATATTGAGAGAGAGTGCCTGAATAACAAGCAACAGCTCATGAGAGCTCGAGAAGCTGCAATTTG
GGAGCTCGAAGAACGACACTTACAAGAAAAACACCAGCTGCTCAAACAGCAGCTTAAAGATCAGTATTTCATGCAAAGACATCAGCTAC
TTAAGCGCCACGAGAAGGAAACAGAGCAAATGCAGCGTTACAATCAAAGACTTATTGAGGAATTGAAAAACAGACAGACTCAAGAAAGA
GCAAGACTGCCCAAGATTCAGCGCAGTGAAGCCAAGACTCGAATGGCCATGTTTAAGAAGAGTTTGAGAATTAACTCAACAGCCACACC
AGATCAGGACCGTGATAAAATTAAACAGTTTGCTGCACAAGAAGAAAAGAGGCAGAAAAATGAGAGAATGGCTCAGCATCAGAAACATG
AGAATCAAATGCGAGATCTTCAGTTGCAGTGTGAAGCCAATGTCCGCGAACTGCATCAGCTGCAGAATGAAAAATGCCACTTGTTGGTT
GAGCATGAGACTCAGAAACTGAAGGAGTTAGATGAGGAACATAGCCAAGAATTAAAGGAGTGGAGAGAGAAATTGAGACCTAGGAAAAA
GACACTGGAAGAAGAGTTTGCCAGGAAACTACAGGAACAGGAAGTATTCTTTAAAATGACTGGGGAGTCTGAATGCCTTAACCCATCAA
CACAGAGCCGGATTTCCAAATTTTATCCTATTCCCAGCTTGCATTCCACCGGATCAGGCAGCGGCAGCGGCAGCGACTACAAAGACCAT
GACGGTGATTATAAAGATCATGATATCGATTACAAGGATGACGATGACAAGGGCAGCGGCAGCGGCAGCGGTAAGCCTATCCCTAACCC
TCTCCTCGGTCTCGATTCTACGTAA

MSFFNFRKIFKLGSEKKKKQYEHVKRDLNPEDFWEIIGELGDGAFGKVYKAQNKETSVLAAAKVIDTKSEEELEDYMVEIDILASCDHP
NIVKLLDAFYYENNLWILIEFCAGGAVDAVMLELERPLTESQIQVVCKQTLDALNYLHDNKIIHRDLKAGNILFTLDGDIKLADFGVSA
KNTRTIQRRDSFIGTPYWMAPEVVMCETSKDRPYDYKADVWSLGITLIEMAEIEPPHHELNPMRVLLKIAKSEPPTLAQPSRWSSNFKD
FLKKCLEKNVDARWTTSQLLQHPFVTVDSNKPIRELIAEAKAEVTEEVEDGKEEDEEEETENSLPIPASKRASSDLSIASSEEDKLSQN
ACILESVSEKTERSNSEDKLNSKILNEKPTTDEPEKAVEDINEHITDAQLEAMTELHDRTAVIKENEREKRPKLENLPDTEDQETVDIN
SVSEGKENNIMITLETNIEHNLKSEEEKDQEKQQMFENKLIKSEEIKDTILQTVDLVSQETGEKEANIQAVDSEVGLTKEDTQEKLGED
DKTQKDVISNTSDVIGTCEAADVAQKVDEDSAEDTQSNDGKEVVEVGQKLINKPMVGPEAGGTKEVPIKEIVEMNEIEEGKNKEQAINS
SENIMDINEEPGTTEGEEITESSSTEEMEVRSVVADTDQKALGSEVQDASKVTTQIDKEKKEIPVSIKKEPEVTVVSQPTEPQPVLIPS
ININSDSGENKEEIGSLSKTETILPPESENPKENDNDSGTGSTADTSSIDLNLSISSFLSKTKDSGSISLQETRRQKKTLKKTRKFIVD
GVEVSVTTSKIVTDSDSKTEELRFLRRQELRELRFLQKEEQRAQQQLNSKLQQQREQIFRRFEQEMMSKKRQYDQEIENLEKQQKQTIE
RLEQEHTNRLRDEAKRIKGEQEKELSKFQNMLKNRKKEVINEVEKAPKELRKELMKRRKEELAQSQHAQEQEFVQKQQQELDGSLKKII
QQQKAELANIERECLNNKQQLMRAREAAIWELEERHLQEKHQLLKQQLKDQYFMQRHQLLKRHEKETEQMQRYNQRLIEELKNRQTQER
ARLPKIQRSEAKTRMAMFKKSLRINSTATPDQDRDKIKQFAAQEEKRQKNERMAQHQKHENQMRDLQLQCEANVRELHQLQNEKCHLLV
EHETQKLKELDEEHSQELKEWREKLRPRKKTLEEEFARKLQEQEVFFKMTGESECLNPSTQSRISKFYPIPSLHSTGSGSGSGSDYKDH
DGDYKDHDIDYKDDDDKGSGSGSGKPIPNPLLGLDST*

#### Supplementary Tables

Table 1 — Primers used in this study

| Target Gene | Forward Primer (5’-3’) | Reverse Primer (5’-3’) | Experiment |
|---|---|---|---|
| GAPDH | AATCCCATCACCATCTTCCAG | CCTTCTCCATGGTGGTGAAGAC | RT-qPCR |
| b-Actin | CAGCCATGTACGTTGCTATC | CTTCATGAGGTAGTCAGTCA | RT-qPCR |
| MYC | AATGAAAAGGCCCCCAAGGTAG | GTCGTTTCCGCAACAAGTCCT | RT-qPCR |
| SNAI1 | CTGCAGGACTCTAATCCAGAGTTTA | CCACTGTCCTCATCTGACAGG | RT-qPCR |
| SNAI2 | ATCTGCGGCAAGGCGTTTTCCA | GAGCCCTCAGATTTGACCTGTC | RT-qPCR |
| ZEB1 | GCACCTGAAGAGGACCAGAG | GTGTAACTGCACAGGGAGCA | RT-qPCR |
| ZEB2 | TTCCTGGGCTACGACCATAC | GCCTTGAGTGCTCGATAAGG | RT-qPCR |
| TWIST1 | CTCAAGAGGTCGTGCCAATC | CCCAGTATTTTTATTTCTAAAGGTGTT | RT-qPCR |
| TWIST2 | GGGAGTGAGCACATTAGCAA | GGGCATGAGTACCCTTAGGA | RT-qPCR |
| CDH1 | TGCCCAGAAAATGAAAAAGG | GTGTATGTGGCAATGCGTTC | RT-qPCR |
| CDH2 | ACAGTGGCCACCTACAAAGG | CCGAGATGGGGTTGATAATG | RT-qPCR |
| CDH3 | AAGATCTTCCCATCCAAACG | CTACAGCGAAGACACCCTCA | RT-qPCR |
| VIM | GCGTGACGTACGTCAGCAATATGA | GTTCCAGGGACTCATTGGTTCCTT | RT-qPCR |
| LAD1 | CTTGAGGCTCTCAGGGGTTG | CTGAGTCCTCTCGGTTGCAG | RT-qPCR |
| Total SLK | GATTGCAGAGGCGAAGGCTG | TCCCGAAGTTCCTGACGTCTAAG | RT-qPCR |
| SLK exon 13 #1 | AGCACCCAAAGAGCTGAGAA | CTGAGCATGCTGGCTTTGTG | RT-qPCR |
| SLK exon 13 #2 | TGAAGTGGAGAAAGCACCCA | TTGTGCAAGCTCCTCTTTCCT | RT-qPCR |
| SLK exon 13 #3 | AAGCACCCAAAGAGCTGAGA | GCATGCTGGCTTTGTGCAAG | RT-qPCR |

| | | | |
|---|---|---|---|
| SLK exon 13 #4 | AGCACCCAAAGAGCTGAGAAA | GAGCATGCTGGCTTTGTGC | RT-qPCR |
| SLK-Short-3xFLAG-V5 | GAACAAGAGTTTGTTCAGAAACAACAGCAAG | CTCCTTCTTTCGGTTCTTCAGCATATTCT | Cloning |
| hSLK_sgRNA | CACCGGAGTATCAGCTAAAAACACG | AAACCGTGTTTTTAGCTGATACTCC | CRISPR knockout of SLK |

Table 2 — Antibodies used in this study

| **Target** | **Supplier, Cat #** | **Experiment (dilution)** |
|---|---|---|
| Anti-FLAG | CST, 8146S | Western blot (1:1,000 in TBST), IF (1:100 in PBS + 1% BSA) |
| Anti-HA | CST, 3724S | Western blot (1:1,000 in TBST), IF (1:100 in PBS + 1% BSA) |
| SLK | Proteintech, 19743-1-AP | Western blot (1:500 in TBST), IF (1:200 in PBS + 1% BSA), Co-IP (3-5µg), MolBoolean (1:200 in Complete Diluent) |
| V5-tag | Invitrogen, R960-25 | Western blot (1:2,000 in TBST), IF (1:100 in PBS + 1% BSA) |
| V5-tag | Novus, NB600-381 | Co-IP (2µg) |
| c-Myc | Abcam, ab32072 | Western blot (1:1,000 in TBST), IF (1:100 in PBS + 1% BSA), Co-IP (2µg) |
| c-Myc | Origene, TA500003 | MolBoolean (1:100 in Complete Diluent) |
| β-Actin | CST, 3700S | Western blot (1:1,000 in TBST), IF (1:750 in PBS + 1% BSA) |
| Vinculin | Invitrogen, 14-9777-82 | Western blot (1:1,000 in TBST) |
| LC3 A/B | CST, 12741S | Western blot (1:1,000 in TBST), IF (1:100 in PBS + 1% BSA) |
| ATM | CST, 2873S | Western blot (1:1,000 in TBST) |
| SQSTM1/p62 | CST, 5114T | Western blot (1:1,000 in TBST) |
| Phospho-SQSTM1/p62 (Thr269/Ser272) | CST, 13121S | Western blot (1:1,000 in TBST) |
| Phospho-ATM (Ser1981) | CST, 13050S | Western blot (1:1,000 in TBST) |
| MX1 | CST, 37849S | Western blot (1:1,000 in TBST) |

| | | |
|---|---|---|
| OAS2 | CST, 54155S | Western blot (1:1,000 in TBST) |
| EpCAM | Invitrogen, 14-9326-82 | Flow Cytometry (0.25 µg/test) |
| Normal Mouse IgG | CST, 68860L | Co-IP (2µg) |
| Normal Rabbit IgG | CST, 2729S | Co-IP (3-5µg) |
| Streptavidin-HRP | Thermo Scientific, N100 | Western blot (1:10,000 in TBST) |
| IRDye 680RD Goat anti-Mouse IgG Secondary Antibody | Li-Cor, 926-68070 | Western blot (1:10,000 in TBST) |
| IRDye® 800CW Goat anti-Rabbit IgG Secondary Antibody | Li-Cor, 926-32211 | Western blot (1:10,000 in TBST) |
| Anti-mouse IgG HRP Conjugate | Promega, W402B | Western blot (1:10,000 in TBST) |
| Anti-rabbit IgG HRP Conjugate | Promega, W401B | Western blot (1:10,000 in TBST) |
| Goat Anti-Mouse IgG H&L (Alexa Fluor® 488) | Abcam, ab150113 | IF (1:1,000 in PBS + 1% BSA) |
| Goat Anti-Rabbit IgG H&L (Alexa Fluor® 555) | Abcam, ab150078 | IF (1:1,000 in PBS + 1% BSA) |
| Goat Anti-Mouse IgG H&L (Alexa Fluor® 555) | Abcam, ab150114 | IF (1:1,000 in PBS + 1% BSA) |
| Goat Anti-Rabbit IgG H&L (Alexa Fluor® 647) | Abcam, ab150079 | IF (1:1,000 in PBS + 1% BSA) |
| Goat Anti-Mouse IgG H&L (Alexa Fluor® 647) | Abcam, ab150115 | IF (1:1,000 in PBS + 1% BSA) |
| Clean-Blot™ IP Detection Reagent (HRP) | Thermo Scientific, 21230 | Western blot (1:1,000 in TBST) |
